# Structural basis for Retriever-SNX17 assembly and endosomal sorting

**DOI:** 10.1101/2024.03.12.584676

**Authors:** Amika Singla, Daniel J. Boesch, Ho Yee Joyce Fung, Chigozie Ngoka, Avery S. Enriquez, Ran Song, Daniel A. Kramer, Yan Han, Puneet Juneja, Daniel D. Billadeau, Xiaochen Bai, Zhe Chen, Emre E. Turer, Ezra Burstein, Baoyu Chen

## Abstract

During endosomal recycling, Sorting Nexin 17 (SNX17) facilitates the transport of numerous membrane cargo proteins by tethering them to the Retriever complex. Despite its importance, the mechanisms underlying this interaction have remained elusive. Here, we report the structure of the Retriever-SNX17 complex determined using cryogenic electron microscopy (cryo-EM). Our structure reveals that the C-terminal tail of SNX17 engages with a highly conserved interface between the VPS35L and VPS26C subunits of Retriever. Through comprehensive biochemical, cellular, and proteomic analyses, we demonstrate that disrupting this interface impairs the Retriever-SNX17 interaction, subsequently affecting the recycling of SNX17-dependent cargos and altering the composition of the plasma membrane proteome. Intriguingly, we find that the SNX17-binding pocket on Retriever can be utilized by other ligands that share a consensus acidic C-terminal tail motif. By showing how SNX17 is linked to Retriever, our findings uncover a fundamental mechanism underlying endosomal trafficking of critical cargo proteins and reveal a mechanism by which Retriever can engage with other regulatory factors.

## Introduction

Plasma membrane (PM) proteins undergo frequent internalization into the endosomal compartment, where they are either routed back to the cell surface for reuse or to lysosomes for degradation. The maintenance of this trafficking process is vital for cellular homeostasis and involves intricate regulatory systems. Among these, the trimeric protein complex Retriever plays a crucial role in identifying PM proteins, also called cargoes, for recycling from endosomes. Composed of VPS35L, VPS26C, and VPS29 (Fig. 1a), Retriever is remotely related to the well-studied endosomal recycling complex Retromer^1–3^, which handles a separate subset of cargoes. Recent studies have revealed that while Retriever shares a similar overall architecture with Retromer, it possesses distinct structural features and regulatory mechanisms^4–7^.

**Fig. 1.**
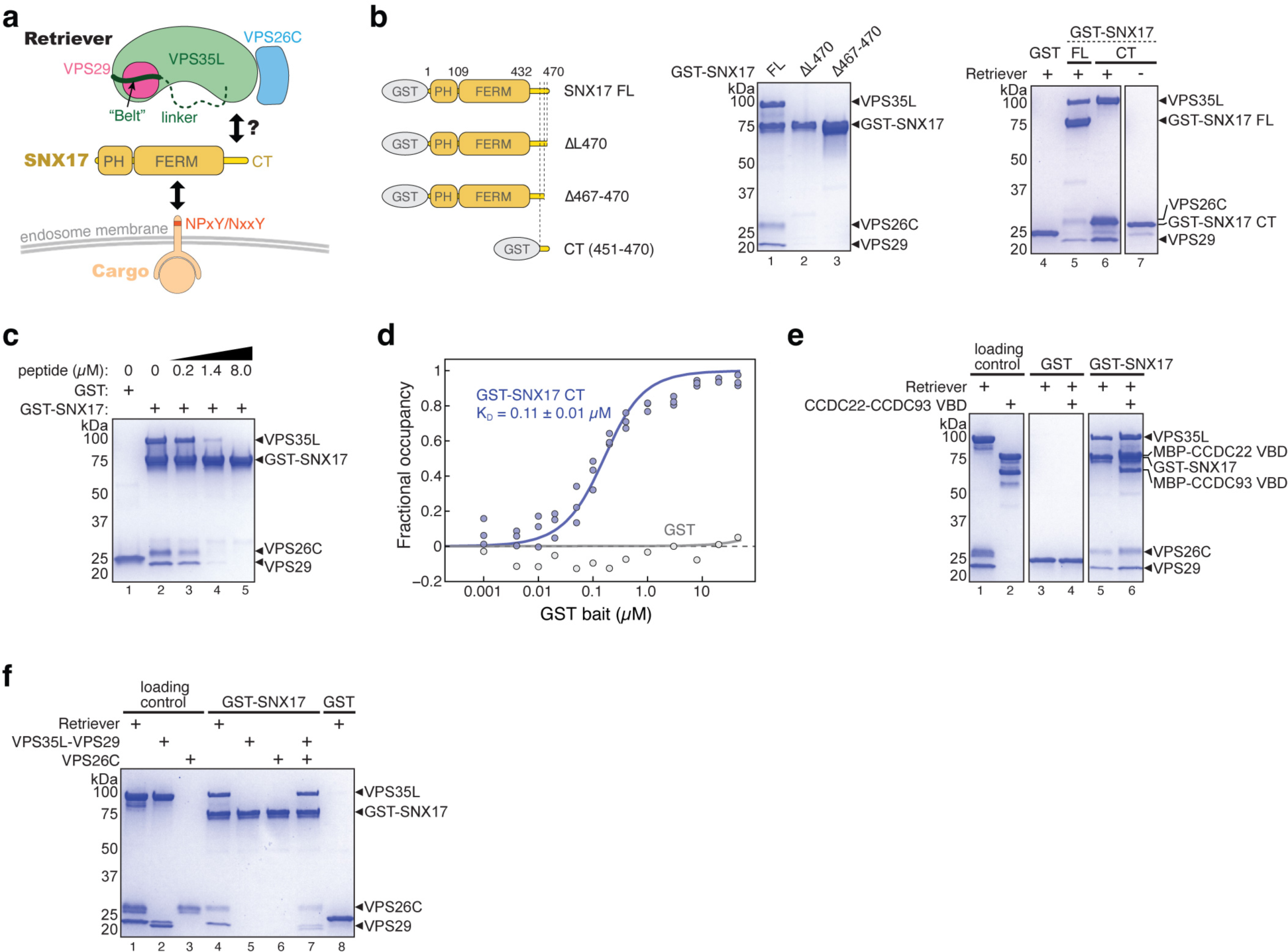
SNX17 uses its C-terminal tail to bind Retriever. **a.** Cartoon depiction of Retriever and the domain architecture of SNX17. **b.** Cartoon representation of GST-SNX17 constructs used (left panel) and Coomassie blue-stained SDS PAGE gels showing in vitro GST pull-down between indicated GST-SNX17 constructs and Retriever (right panel). **c.** Coomassie blue-stained SDS PAGE gels showing in vitro GST pull-down between GST-SNX17 and Retriever in the presence of increasing concentrations of a competing peptide consisting of the last 20 amino acids of SNX17. **d.** Binding isotherms obtained from EPD assays measuring the binding affinity between GST-SNX17 CT and Retriever. Data were pooled from three independent experiments and globally fitted to a one-binding site model to obtain the K_D_ and fitting error^32^. GST pull-down as a negative control was from one experiment. Representative Coomassie blue-stained SDS-PAGE gels from the EPD experiments are shown in Extended Data Fig. 1a. **e-f.** Coomassie blue-stained SDS PAGE gels showing in vitro GST pull-down between GST-SNX17, and Retriever complexed with CCDC22-CCDC93 VBD dimer **(e)** or isolated subunits of Retriever **(f)**. Representative results from at least two independent experiments are shown.

Retriever manages the recycling of a broad spectrum of cargoes, including integrins, tyrosine receptor kinases, G-protein coupled receptors (GPCRs), and lipoprotein receptors^4,8,9^. In contrast, Retromer handles a distinct subset of cargoes, which includes various transporters (DMT1, ATP7A/B, GLUT1) and SorL1 (a sorting factor implicated in Alzheimer’s disease)^10–14^. Both Retriever and Retromer cooperate with additional factors to ensure efficient cargo sorting. Integral to the function of both complexes is the WASH regulatory complex, which promotes Arp2/3-mediated actin polymerization at endosomal membranes^15–18^. In addition, Retriever associates with the COMMD/CCDC22/CCDC93 complex (CCC)^8^ to form a larger-order complex known as the Commander assembly^19–21^, in which the ten COMMD proteins form a ring-like structure^5–7^, while the CCDC22-CCDC93 dimer uses different domains to bridge the COMMD ring to Retriever while also interacting with DENND10, a putative Rab guanine nucleotide exchange factor (GEF).

Sorting nexin proteins represent crucial regulatory factors responsible for tethering Retromer or Retriever to endosomal membranes and their specific cargoes^22^. Retromer-associated sorting nexins, like SNX1, SNX2, SNX5 and SNX6, mediate membrane deformation, while SNX3 and SNX27 tether Retromer to particular cargoes. For example, SNX27 links Retromer to over 100 cargo proteins by simultaneously binding to the VPS26 subunit of Retromer and the PDZ binding motif in the cytoplasmic tail of the cargoes. In contrast, SNX17, a distant homolog of SNX27, is specifically associated with Retriever. Unlike SNX27, SNX17 uses its protein 4.1R, ezrin, radixin, moesin (FERM) domain to recognize the NPxY/NxxY motif in the cytoplasmic tail of over 100 distinct cargo proteins^23,24^, while its interaction with Retriever involves its C-terminal tail^4^. The precise mechanism underlying the Retriever-SNX17 interaction, however, has yet to be deciphered (Fig. 1a). It also remains unknown if other regulatory factors connect Retriever to additional cargos or recycling processes.

Here, we present the cryo-EM structure of the Retriever-SNX17 complex and comprehensive validation of the binding mechanism through biochemical, cellular, and proteomic analyses. Furthermore, we report the discovery of additional ligands for Retriever, which similarly interact with the complex through the conserved SNX17-binding pocket. This finding expands the repertoire of regulatory factors of Retriever and suggests versatile connections of Retriever with other potential recycling targets.

## Results

### SNX17 uses its C-terminal tail to bind Retriever

Previous cellular and co-immunoprecipitation studies showed that the C-terminal (CT) unstructured tail of SNX17 is important for interacting with Retriever^4^ (Fig. 1a). Here, we used recombinantly purified proteins to determine whether the interaction is direct and further elucidate how the interaction occurs. Our GST pull-down assays showed that GST-SNX17 directly interacted with Retriever and, consistent with previous cell-based results^4^, the in vitro interaction relied on the C-terminal tail of SNX17 (Fig. 1b). Deleting the last four residues (Δ467-470) or the last residue (Δ470) of the tail abolished the interaction (Fig. 1b, lanes 2-3). We found that the tail was both necessary and sufficient for the interaction, as a GST-tagged tail peptide, comprising the last 20 residues, similarly pulled down Retriever (Fig. 1b, lane 6), and a chemically synthesized peptide of the same 20 residues of the tail could compete off the binding of GST-SNX17 in a dose-dependent manner (Fig. 1c). Using an equilibrium pull-down assay, we determined that the binding has a dissociation constant (K_D_) of ∼0.11 µM in our buffer condition (Fig. 1d; Extended Data Fig. 1a). In addition, in the same in vitro conditions, we found that SNX17 could similarly bind to Retriever complexed with the VPS35L binding domain (VBD) of the CCDC22-CCDC93 dimer (Fig 1e, lane 6), the key scaffold required for CCC complex assembly^6^, suggesting that SNX17 interacts similarly with Retriever alone or with the Retriever-CCC complex.

Intriguingly, SNX17 could not bind individual subunits of Retriever, including a VPS35L-VPS29 binary subcomplex or the VPS26C subunit in isolation, and only bound to fully assembled Retriever (Fig. 1f, lanes 4-6). This is not due to misfolding or mis-assembly of the isolated components, as the interaction was readily recovered when the individually purified VPS35L-VPS29 subcomplex and VPS26C were freshly mixed in the reaction (Fig. 1f, lane 7; also see Extended Data Fig. 1b for size exclusion chromatography profiles of individual components indicating monodispersed, well-behaving materials). The above results confirm the requirement of VPS26C for binding^4^ and suggest that SNX17 only directly interacts with fully assembled Retriever in vivo.

### Cryo-EM structure of Retriever-SNX17 complex

To understand how the SNX17 tail interacts with Retriever, we next determined the structure of the Retriever-SNX17 complex using cryo-EM. After exhaustively surveying protein constructs and grid conditions, we were able to obtain a cryo-EM map with a resolution of ∼3.4 Å by using Retriever mixed with saturating concentrations of the SNX17 tail peptide (Fig. 2a, Table 1, Extended Data Fig. 2). We used local refinement and local resolution-based map sharpening^25^ to improve map quality and built the structural model starting with one generated by AlphaFold-Multimer prediction (Fig. 2a and Extended Data Fig. 2 & 3a). The overall crescent-shaped structure of Retriever is slightly extended compared to its apo form, with an average root-mean-square deviation of ∼1.9 Å (Extended Data Fig. 3b). Due to potential conformational dynamics, we could not obtain a well-resolved map for the VPS29-bound end, where the N-terminal “belt” sequence of VPS35L was found to stabilize the bound VPS29 and the CT region of VPS35L in our previous work^6^ (Fig. 2a, represented by dashed line, and Extended Data Fig. 2 and 3b).

**Fig. 2.**
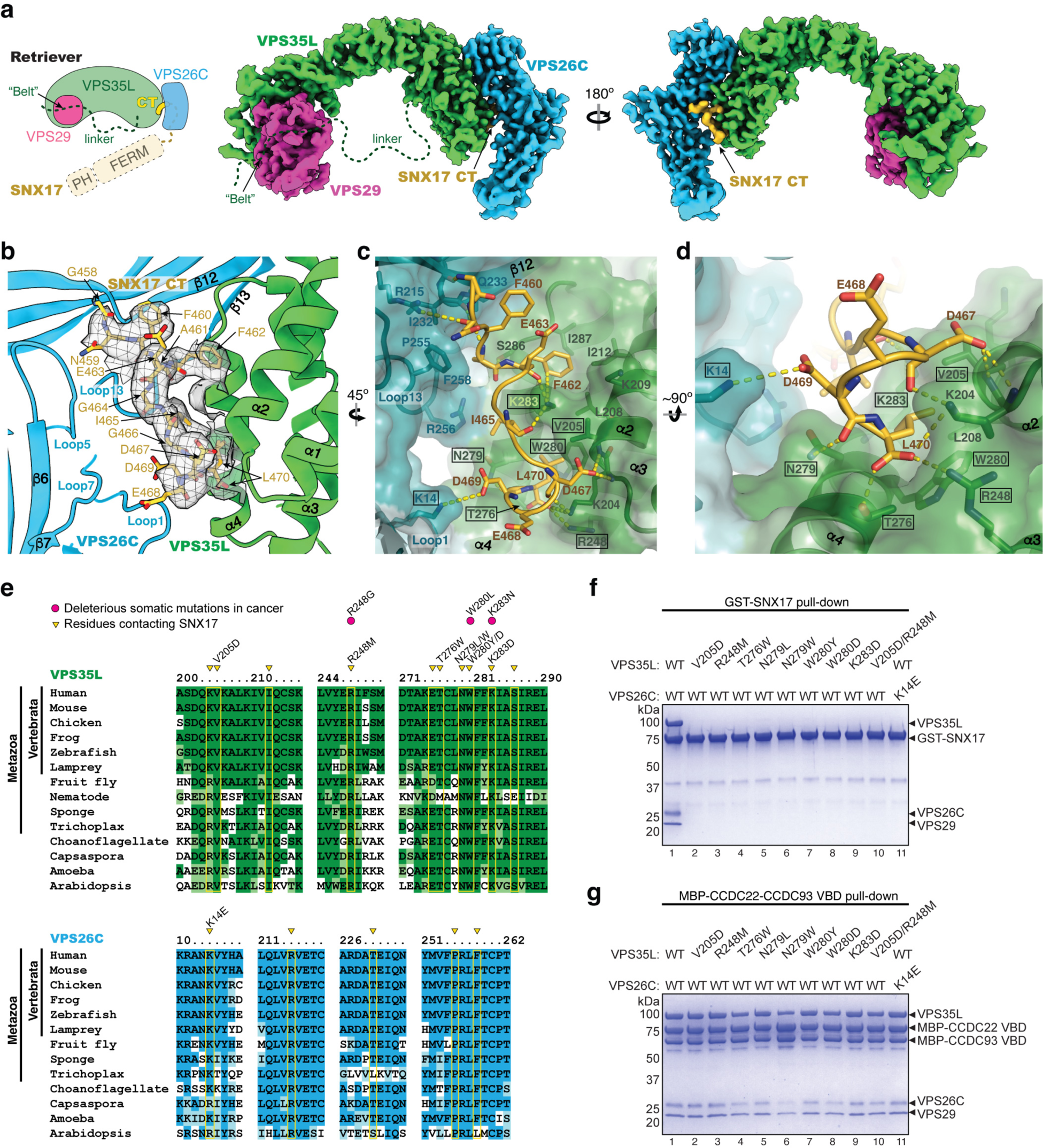
Structure of SNX17 C-terminal tail binding to Retriever. **a.** Schematic and overall colored cryo-EM map of the SNX17 CT peptide (golden) complexed with Retriever (VPS35L in green, VPS29 in magenta, and VPS26C in cyan). Dotted lines indicate structural elements not resolved in cryo-EM. N-terminal domains of SNX17 are shown as a reference. **b-d.** Close-up views showing key interactions between the SNX17 CT peptide (carbon in green, oxygen in red, and nitrogen in blue) and its binding surface on VPS35L and VPS26C: **(b)** shows cryo-EM density of the SNX17 CT peptide; **(c-d)** shows surface conservation calculated with ConSurf^33^, with color-to-white gradients representing the most (ConSurf score = 9) to the least conserved residues (ConSurf score = 1). Contacting residues are shown as sticks. Dotted yellow lines indicate polar interactions. Residues mutated in this study are indicated by a black box. **e.** Sequence alignment of human VPS35L and VPS26C with orthologs from indicated representative species. Residues contacting SNX17 are indicated with yellow boxes and arrowheads. Deleterious somatic mutations found in the COSMIC database and mutations tested for binding to SNX17 are indicated. **f-g.** Coomassie blue-stained SDS PAGE gels showing GST-SNX17 **(f)** or MBP-CCDC22-CCDC93 VBD dimer **(g)** pulling down purified Retriever bearing the indicated point mutations in VPS35L or VPS26C. Representative results from at least two independent experiments are shown.

**Table 1.** Cryo-EM data collection and model statistics.

|  | VPS35L-VPS29-VPS26C-SNX17<br>(EMD-43873) | VPS35L<br>(partial)-<br>VPS26C-SNX17<br>(EMD-43870) | VPS35L<br>(partial)-<br>VPS29<br>(EMD-43871) | Composite map<br>(EMD-43872)<br>(PDB 9AU7) |
| --- | --- | --- | --- | --- |
| Data collection and processing |  |  |  |  |
| Magnification | 105,000 |  |  |  |
| Voltage (kV) | 300 |  |  |  |
| Electron exposure (e<sup>-</sup>/Å<sup>2</sup>) | 64 |  |  |  |
| Defocus range (μm) | -0.9 to -2.4 |  |  |  |
| Pixel size (Å) | 0.83 |  |  |  |
| Symmetry imposed | C1 |  |  |  |
| Initial particle images (no.) | 1,009,886 |  |  |  |
| Final particle images (no.) | 227,973 |  |  |  |
| Map resolution (Å) | 3.4 | 3.35 | 3.75 |  |
| FSC threshold | 0.143 |  |  |  |
| Refinement |  |  |  |  |
| Initial model used |  |  |  | AlphaFold<br>Multimer model<br>(ma-swt4h) |
| Model composition |  |  |  |  |
| Nonhydrogen atoms |  |  |  | 8601 |
| Protein residues |  |  |  | 1079 |
| Ligand |  |  |  | 0 |
| R.m.s. deviations |  |  |  |  |
| Bond lengths (Å) |  |  |  | 0.005 |
| Bond angles (°) |  |  |  | 1.026 |
| Validation |  |  |  |  |
| MolProbity score |  |  |  | 1.93 |
| Clashscore |  |  |  | 13.06 |
| Poor rotamers (%) |  |  |  | 0.21 |
| Ramachandran plot |  |  |  |  |
| Favored (%) |  |  |  | 95.65 |
| Allowed (%) |  |  |  | 4.35 |
| Disallowed (%) |  |  |  | 0.00 |
| Protein residues included in the model |  |  |  | VPS35L:115-131, 180-253, 264-349, 352-738, 742-755, 768-786<br>VPS29: 3-186<br>VPS26C: 1-53, 63-146, 150-297<br>SNX17: 458-470 |

Nevertheless, the map unambiguously located the density of SNX17’s CT tail over a conserved surface nestled between the VPS35L and VPS26C subunits of Retriever (Fig. 2a, b, with map quality shown in Extended Data Fig. 3c, d). This density readily accommodated 12 residues at the C-terminus of the peptide (Fig. 2a, b). The peptide adopts a uniquely twisted conformation containing two short, distorted 1-turn helices separated by a short loop (Fig. 2c, d). The majority of the interaction is mediated by a conserved and positively charged surface on VPS35L, contributed largely by residues from helices α2, α3, and α4 (Fig. 2b, c; Extended Data Fig. 4 a, b). In addition, a conserved and slightly positively charged surface on VPS26C, mainly contributed by residues on Loop 1, Loop 13, and strand β12, interacts with the SNX17 peptide from the opposite side (Fig. 2c; Extended Data Fig. 4a, b). This interaction is unique to Retriever, as Retromer uses distinct surfaces to interact with SNX3, SNX27, or DMT1^26–28^ (Extended Data Fig. 4c). This experimentally derived cryo-EM model of the Retriever-SNX17 complex is consistent with AlphaFold-Multimer predictions, with some differences in the residues leading to the C-terminal tip of SNX17 (Extended Data Fig. 3a).

The structural model elucidates the significance of the last few amino acids of SNX17 previously shown to be critical to the binding to Retriever^4^. Remarkably, the last residue of SNX17, L470, uses both its side chain and the carboxyl group to establish a network of interactions with VPS35L critical for binding (Fig. 2c, d). Specifically, L470’s carboxyl group engages with residues K204, R248, and T276 in VPS35L, while its side chain fits into a deep hydrophobic pocket formed by V205, W280, and the alkyl chains of K204 and K283 of VPS35L. These interactions explain why mutating L470 to G or deleting this residue abolished the Retriever-SNX17 interaction^4^ (Fig. 1b, lane 2).

In addition to L470, the structure also explains how other residues in SNX17’s tail contribute to the binding (Fig. 2c, d). At the C-terminal portion of the peptide, D469 in SNX17 is oriented towards K14 from VPS26C, while D467 in SNX17 and K204 in VPS35L engage with each other’s backbone. At the N-terminal portion of the peptide, F462 in SNX17 interacts with residues L208, I212, I287, and K283 in VPS35L, with K283 in VPS35L further interacting with the backbone of F462 and I465 in SNX17. In addition, residues N459, F460, and A461 of SNX17 may form van der Waals interactions with the VPS26C surface.

It is remarkable that the sequences of both VPS35L and VPS26C that contribute to the SNX17 binding pocket, especially the residues directly involved in the interaction, are conserved across a diverse range of organisms from human to amoeba and *Arabidopsis* (Fig. 2c-e; Extended Data Fig. 4a). This suggests that the SNX17-Retriever interaction mechanism is conserved through evolution. Moreover, at least three of these SNX17-interacting residues have been noted to be mutated in cancer (Fig. 2e, indicated by pink dots)^6^, with the resulting missense change predicted to be deleterious.

To validate our structural model, we purified a series of Retriever complexes in which we mutated individual residues that make critical contacts with the SNX17 tail, either from the VPS35L or VPS26C side. We then used GST pull-down experiments to examine how these mutations affect the SNX17-Retriever interaction. Consistent with our structure, all mutations abolished the binding to GST-SNX17 (Fig. 2f). Importantly, the disruption of the binding was not due to misassembly of Retriever, as the mutant complexes behaved similarly to their wild-type (WT) counterparts during protein expression, purification, and size-exclusion chromatography (Extended Data Fig. 1b, c). Furthermore, the mutations did not affect the binding of Retriever to the CCDC22/CCDC93 VPS35L binding domain (VBD) dimer (Fig. 2g; Extended Data Fig. 1d), which is mediated by different conserved surfaces on the VPS29-bound end of the complex, away from the SNX17 binding pocket (Extended Data Fig. 4a), further supporting that the mutations were specific in disrupting the binding to SNX17. Thus, we postulate that the identified SNX17-binding pocket (hereafter named SBP) is evolutionarily conserved and plays a specific role in binding to SNX17.

### Disrupting the SBP alters SNX17 binding in cells

Having defined the SBP as required for in vitro binding between Retriever and SNX17, we examined whether this interaction mechanism held true in cells using co-immunoprecipitation experiments. First, we observed that in transfected HEK293T cells, different mutations in the SBP impacted the binding between SNX17 and Retriever to varied extents. The mutations V205D and R248M in VPS35L substantially weakened the interaction, while other mutations (N279L and W280Y) had minimal effects (Fig. 3a). Combining the V205D and R248M mutations (denoted as DM for double mutant hereafter) had a more profound impact on SNX17-Retriever binding (Fig. 3b). Next, using immunoprecipitation in the reciprocal direction further confirmed the significant contribution of VPS35L SBP residues (N279, W280, V205, and R248) to the interaction between Retriever and SNX17 (Fig. 3c), with VPS35L DM displaying the most robust impairment. Importantly, all the mutants tested retained normal interactions in cells with the Retriever subunits VPS29 and VPS26C, as well as normal interactions with the CCC complex (CCDC22 and DENND10) (Fig. 3c), confirming our in vitro results that the mutations specifically disrupted Retriever binding to SNX17 without affecting other regions of Retriever.

**Fig. 3.**
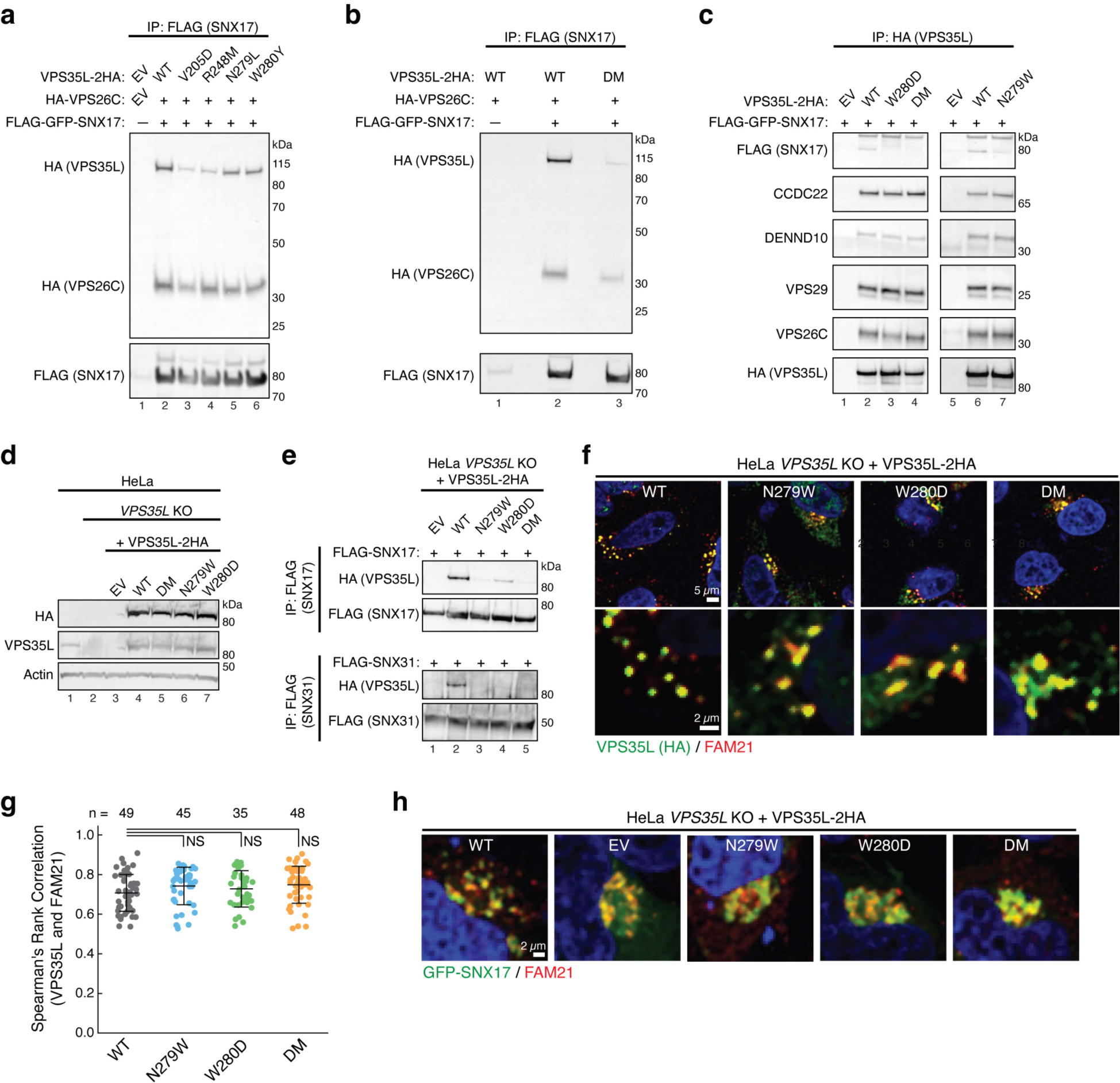
Disrupting the SBP impairs SNX17 and SNX31 binding in cells. **a-b.** Immunoprecipitation of SNX17 (FLAG) followed by immunoblotting for VPS35L and VPS26C (HA) in HEK293T cells transfected with the indicated expression vectors. EV, empty vector; DM, double mutant. **c.** Immunoprecipitation of VPS35L (HA) followed by immunoblotting for SNX17 (FLAG) and indicated protein components of the CCC and Retriever complexes in HEK293T cells transfected with indicated SNX17 and VPS35L variants. **d.** Immunoblotting analysis for endogenous and stably expressed VPS35L in the indicated HeLa cell lines derived from a *VPS35L* knockout (KO) rescued with the indicated variants of VPS35L or an empty vector (EV) control. The parental HeLa cell line used to derive the VPS35L knockout line is included for comparison. **e.** Immunoprecipitation of SNX17 (top) or SNX31 (bottom) after transfection in the indicated HeLa cell lines, followed by immunoblotting for VPS35L (HA). **f-g.** Representative confocal images (**f**) and quantification of colocalization (**g**) derived from concurrent immunofluorescence staining for VPS35L (HA, green) and the endosomal marker FAM21 (red) in HeLa cells shown in (**e**). In (**g**), each dot represents an individual cell, with number of cells in each group indicated above the graph. Mean and standard deviation are shown. One-way ANOVA with Dunnett’s correction was used. NS, not significant. **h.** Representative confocal images showing concurrent immunofluorescence staining for GFP-SNX17 (green) and the endosomal marker FAM21 (red) in HeLa cells shown in (**e**) and transfected with GFP-SNX17.

Finally, we complemented a previously established *VPS35L* knockout (KO) HeLa cell line^8^ and generated polyclonal sublines stably re-expressing different HA-tagged VPS35L variants, including WT, N279W, W280D, and DM, or a control line transfected with an empty vector (EV) (Fig. 3d). Using these lines, we examined whether the SBP is required for SNX17 as well as SNX31 binding. SNX31 is a homolog of SNX17 (40% identity between human proteins) expressed only in very few cell types. SNX31 was previously found to bind to Retriever in a manner that also required its CT leucine residue^4^. Our co-immunoprecipitation experiments demonstrated that both SNX17 and SNX31 bound to VPS35L WT but not to the SBP mutants (Fig. 3e), indicating that the SBP is required for Retriever interactions with both proteins.

### Retriever-SNX17 binding is not required for their endosomal localization

Next, we examined the potential impact of decoupling Retriever from SNX17 on the localization of these proteins in cells. Using immunofluorescence staining in the aforementioned stable lines shown in Fig. 3d, we found that the re-expressed VPS35L was localized to FAM21-positive endosomes regardless of mutations in the SBP (Fig. 3f, g). Reciprocally, SNX17 localization in these cells was assessed after transfection of GFP-SNX17, showing that endosomal localization of SNX17 appeared normal in EV cells and was not impacted by disruption of the SBP (Fig. 3h). Thus, endosomal recruitment of VPS35L and SNX17 are both independent of Retriever-SNX17 complex formation.

### Disruption of the SBP alters PM homeostasis

Next, we examined the functional consequence of disrupting the Retriever-SNX17 interaction on endosomal protein sorting and PM protein homeostasis. To assess this, we first utilized surface biotinylation, protein isolation, and mass spectrometry, coupled with tandem mass tag (TMT) quantification, to compare the PM protein presentation in HeLa cells re-expressing VPS35L WT or the DM mutant, or the knockout line (complemented with EV). This method detected 25 proteins that were significantly reduced in *VPS35L*-deficient cells (EV) after surface biotinylation. These included several integrins, which are known cargoes of SNX17 (Fig. 4a). Consistent with previous observations, several cytosolic proteins were also detected, possibly through indirect interactions with the biotinylated proteome. These included VPS35L itself, which was depleted in the EV cells, as expected. Remarkably, all cargoes reduced in EV cells were similarly reduced in VPS35L DM mutant cells (Fig. 4a), suggesting that the Retriever-SNX17 interaction plays a major role in Retriever-mediated cargo sorting and recycling to the PM.

**Fig. 4.**
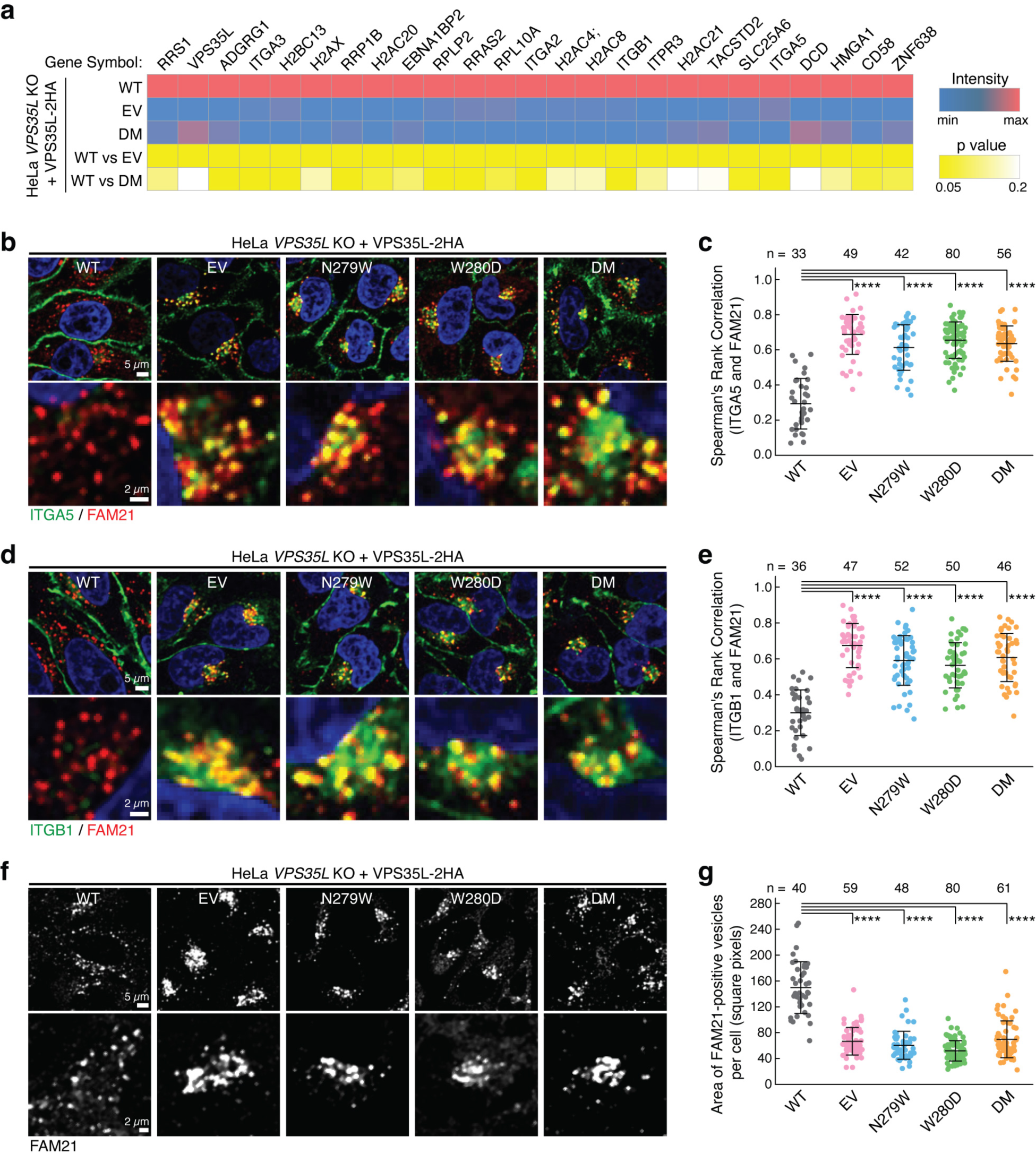
Disrupting the SNX17-Retriever interaction impairs PM homeostasis. **a.** Heat map of PM protein abundance quantified by TMT-based proteomics after surface biotinylation and streptavidin purification, using indicated HeLa stable cell lines (shown in Fig. 3d). Only proteins whose abundance was significantly different between WT and EV lines (p value <0.05) are displayed. **b-c.** Representative confocal images (**b**) and quantification of colocalization (**c**) derived from concurrent immunofluorescence staining for the cargo protein ITGA5 (green) and the endosomal marker FAM21 (red) in the indicated cell lines. **d-e.** Similar to (**b-c**) but focusing on the cargo protein ITGB1 (green). **f-g.** Representative confocal images (**f**) and quantification of the area of FAM21-positive endosomes (**g**) derived from immunofluorescence staining of FAM21 in the indicated HeLa stable cell lines. In all quantifications, each dot represents an individual cell, with number of cells in each group indicated above the graph. Mean and standard deviation are shown. One-way ANOVA with Dunnett’s correction was used. ****, p < 0.0001.

To validate the PM proteomics result, we used immunofluorescence staining to directly evaluate the cellular localization of Integrin α5 (ITGA5). Consistent with the proteomics data, ITGA5 exhibited reduced staining at the PM and significant accumulation in FAM21-positive endosomes of EV cells (Fig. 4b, c). Importantly, SBP mutations in VPS35L led to a comparable phenotype, showing significant endosomal trapping of ITGA5 (Fig. 4b, c). The same analysis of another SNX17 cargo, Integrin β1 (ITGB1), revealed a similar pattern of endosomal trapping in cells lacking VPS35L (EV) or expressing VPS35L with SBP mutations (Fig. d, e). Associated with the endosomal trapping phenotypes was an altered morphology of FAM21-positive endosomes, displaying enlarged endosomal domains and coalescence in the perinuclear region (Fig. 4f). The coalescence phenotype, quantified as area of FAM21-positive endosomes per cell, showed significant alterations in VPS35L deficiency (EV) as well as in all SBP mutants tested (Fig. 4g). Thus, decoupling Retriever from SNX17 had a profound effect on endosomal sorting and homeostasis of various PM proteins.

### SBP mutations reveal other acidic tail partners of Retriever

Next, we assessed the range of protein-protein interactions for VPS35L WT and compared its interacting partners with the DM variant. To accomplish this, we immunoprecipitated VPS35L from the corresponding HeLa cell lines and used mass spectrometry to identify interacting partners in an unbiased manner. Using 10-fold enrichment over the EV knockout as a threshold, coupled with statistical testing, we identified 25 potential VPS35L interacting proteins (Fig. 5a). This analysis readily identified all components of the Retriever and CCC complexes (Fig. 5a, green labels), but could not detect SNX17, potentially due to its cellular abundance, low stoichiometry or affinity of binding, or poor peptide ionization. Our method also identified several proteins not previously reported to be partners of Retriever such as LRMDA, CDIPT, ADGRE3, PFH5A, DYNLL1 RNF169, BAG2 and LRP12 (Fig. 5a, orange labels).

**Fig. 5.**
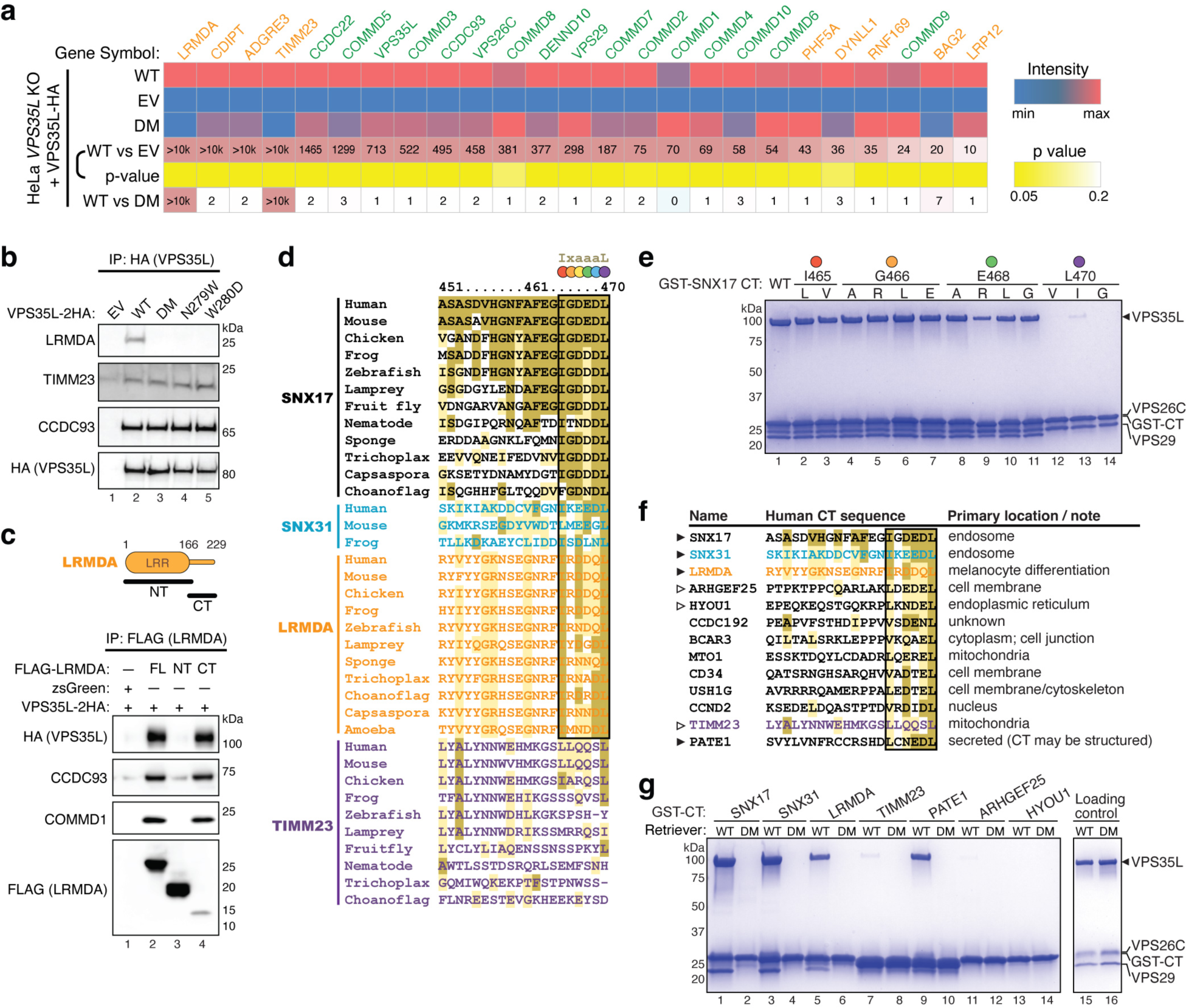
Interactome analysis reveals other Retriever ligands with an acidic CT tail. **a.** Heat map of VPS35L-interacting proteins quantified by mass spectrometry after coimmunoprecipitation of HA-tagged VPS35L, using indicated HeLa stable cell lines shown in Fig. 3d. Protein spectral counts, fold change between indicated cells, and corresponding p-values are depicted. Components of CCC and Retriever are highlighted in green, while other proteins are marked in orange. **b.** Immunoprecipitation of VPS35L (HA) followed by immunoblotting for LRMDA, TIMM23, and CCDC93 in indicated stable HeLa cell lines. **c.** Immunoprecipitation of LRMDA full-length (FL), NT, and CT in Lenti-X 293T cells, followed by immunoblotting for indicated Retriever and CCC subunits. **d.** Sequence alignment of the CT tail of indicated proteins across representative species. Gold, light-gold, and white shading denote sequences identical to, similar to, and not conserved with the human SNX17 CT sequence, respectively. The identified 6-residue acidic sequences are highlighted by the black box, with each position denoted by a colored dot corresponding to the residues mutated in (**e**). **e.** Coomassie blue-stained SDS PAGE gel showing in vitro pull-down of Retriever by GST-SNX17 CT tails containing indicated point mutations. **f.** Sequence alignment of the CT tail of indicated proteins in humans, of which the last six residues concur to the motif [LVI]-X-[DEQN]-X-[DEQN]-L and exist in unstructured tails. Gold, light-gold, and white shading denote sequences identical to, similar to, and not conserved with the human SNX17 CT sequence, respectively. **g.** Coomassie blue-stained SDS PAGE gel showing in vitro pull-down of Retriever WT vs. the DM mutant by GST-tagged CT acidic tails of the indicated proteins.

Intriguingly, we found that two proteins, LRMDA (leucine rich melanocyte differentiation associated) and TIMM23 (translocase of inner mitochondrial membrane 23), preferentially bound to VPS35L WT but not the DM mutant (enrichment ratio of 10 or greater). Immunoprecipitation and Western blot confirmed that LRMDA only interacted with VPS35L WT but not the SBP mutants (Fig. 5b). In contrast, TIMM23 appeared to bind equally to all mutants (Fig. 5b). LRMDA contains an NT leucine-rich repeat (LRR) domain and a CT unstructured tail (Fig. 5c, cartoon). Immunoprecipitation in Lenti-X 293T cells transfected with the full length (FL), NT LRR, and CT tail of LRMDA revealed that the CT tail is both necessary and sufficient for binding to Retriever (Fig. 5c), analogous to SNX17.

Comparing the CT tail sequences of validated SBP-dependent binders (SNX17, SNX31, and LRMDA) versus the non-SBP-dependent binder (TIMM23) across various representative species revealed the presence of significant homology among their extreme C-terminus (Fig. 5d), suggesting a potentially shared mechanism of binding. This homology can be summarized as an evolutionarily conserved consensus motif comprising the last 6 residues, denoted as I-x-aaa-L, where “a” denotes an acidic amino acid residue (Fig. 5d, black box). The sequences preceding this motif lack a discernable pattern, suggesting the decisive role of this motif in binding to Retriever.

In this motif, the last Leu residue seemed the most invariant, followed by the first Ile residue, which could be Leu or Phe in several species. The three central acidic residues could be a combination of Asp, Glu, Asn, and Gln, while the x position appeared to tolerate various amino acids. To further understand the composition of this motif, we performed mutagenesis of various residues in the CT acidic tail of SNX17 and tested how they affected the binding to Retriever (Fig. 5e). We found that any mutations of the terminal Leu abrogated the binding, consistent with the sequence alignment analysis in Fig. 5d. Mutations of the central acidic residue did not disrupt the binding, except the mutation to Arg, which weakened the binding. This is consistent with the cryo-EM structure, in which E468 side chain points away from the binding surface to the solvent (Fig. 2d). Mutating I465 to Leu or Val, or mutating the “x” position (G466) to various amino acids did not have an appreciable effect on binding in vitro (Fig. 5e). Combining the sequence alignment and the mutagenesis of SNX17 tail, we redefined the consensus motif as [I/L/V]-x-[DEQN]-x-[DEQN]-L.

Based on this motif, we searched the human genome and identified additional proteins containing such a sequence in an unstructured C-terminal tail (Fig. 5f, Extended Data Fig. 5). These proteins have diverse membrane localizations and were previously not shown to bind to Retriever. When fused to GST, the last 20 amino acids of SNX17, SNX31, LRMDA, and PATE1 supported robust binding to Retriever in vitro (Fig. 5g), while TIMM23 showed a weak interaction. Notably, all interactions were abrogated by the DM mutation, confirming that they use the same mechanism to interact with the SBP of Retriever (Fig. 5g).

## Discussion

Our study provides a pivotal advance in our understanding of Retriever-mediated endosomal cargo recycling. Unlike the well-studied Retromer^26,28,29^, the precise mechanisms of cargo selection by Retriever have remained elusive^30^. Our findings elucidate how the cargo-recognition factor SNX17 uses its acidic C-terminal tail to anchor into a conserved surface formed by the VPS35L and VPS26C subunits of Retriever (Fig. 6). Furthermore, we have identified other regulatory factors that interact with Retriever through the SBP and via similar acidic tails, suggesting that the SBP is a critical surface that connects Retriever to other cellular functions beyond SNX17-dependent cargo recycling^8^ (Fig. 6).

**Fig. 6.**
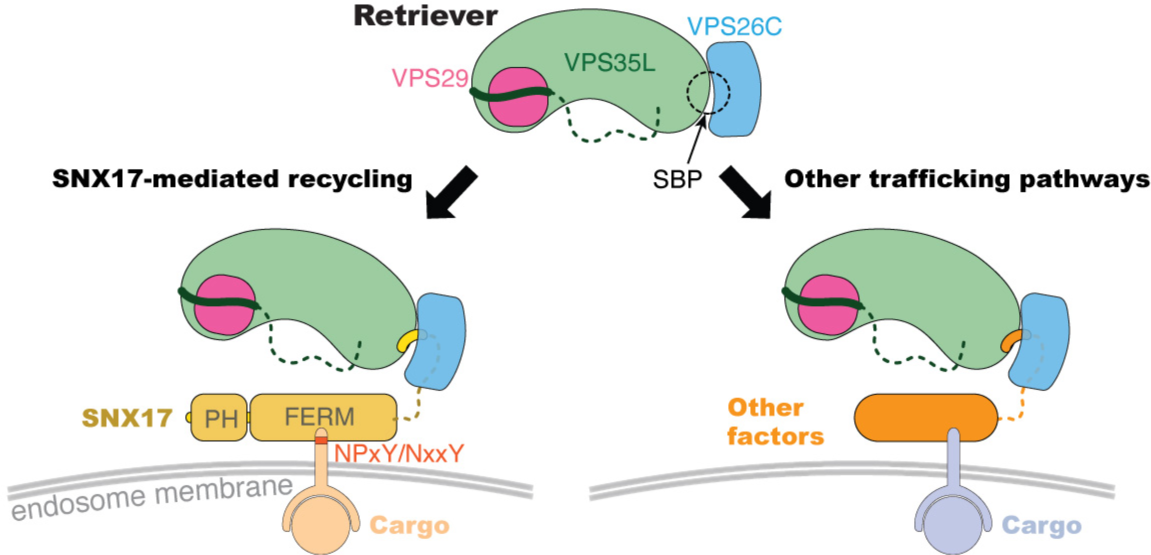
Schematic of Retriever-mediated endosomal recycling. Our study suggests that Retriever could use the same SNX17-binding pocket (SBP) to interact with additional factors. The interaction with SNX17 tethers Retriever to many cargoes recognized by SNX17 through NPxY/NxxY motifs in their cytoplasmic tails (left), while the interaction with other factors can potentially link Retriever to distinct cargoes, recycling pathways, or cellular destinations (right).

We emphasize that despite sharing remote homology with Retromer and SNX27, Retriever and SNX17 operate through very distinct mechanisms. Notably, the residues surrounding the binding pocket on both the VPS35L and VPS26C sides are highly conserved across species, representing one of the most conserved surfaces on Retriever (Extended data Fig. 4a). This remarkable conservation underscores the evolutionary and functional significance of the interaction between Retriever and SNX17, as well as other acidic tail co-factors.

Another striking observation is that disrupting the Retriever-SNX17 interaction has profound consequences on PM protein homeostasis, with effects on cellular signaling and potential clinical implications. This is evidenced by our proteomic and cellular analyses, as well as the association between somatic mutations at the SNX17 binding pocket and human cancers. Mutations at these residues disrupt the Retriever-SNX17 interaction in our experimental system, suggesting that the cancer-associated mutations may act by perturbing the homeostasis of crucial cargoes involved in cell adhesion, proliferation, or metabolism. This binding pocket, therefore, offers a promising target for the development of novel therapeutic interventions or small molecule drugs to modulate cellular signaling dynamics.

Our proteomic and cellular studies indicate that SNX17 does not constitutively bind to Retriever. It is plausible that the binding could be modulated by factors such as SNX17’s expression level, post-translational modifications, cargo density, and cellular localization. For example, based on PhosphoSitePlus^31^, two key residues at the SBP could be potentially modified, including acetylation of K14 in VPS26C and ubiquitylation of K204 in VPS35L, which could disrupt the binding.

Moreover, the identification of other factors containing the SNX17 homologous acidic tail sequences suggests a versatile role for the binding pocket. These additional factors may act as competitors of SNX17 and connect Retriever to a broader range of recycling pathways, cargoes, or other cellular locations and functions. Given the conservation and functional importance of the binding pocket, we speculate that pathogens might exploit this system by hijacking the Retriever-SNX17 interaction with effector proteins that mimic the critical C-terminal motif, thus compromising host cellular functions to create a niche or augment pathogen fitness. In summary, our research not only elucidates a key mechanism in endosomal trafficking but also opens the door for further exploration into the biological significance of other Retriever-ligand interactions.

## Acknowledgements

GST-SNX17 FL construct was a gift from Titus Boggon at Yale University School of Medicine. We thank the Research IT at Iowa State University for hardware resources, installation of AlphaFold, and ongoing computational & diagnostic support. We also thank Andrew Lemoff and the Proteomics core at UT Southwestern. Electron Microscopy data were collected in collaboration with the Structural Biology Laboratory using the Cryo Electron Microscopy Facility at UT Southwestern Medical Center (partially supported by grant RP220582 from the Cancer Prevention & Research Institute of Texas [CPRIT] for cryo-EM studies) and the Iowa State University Cryo-EM Facility (supported by the Roy J. Carver Structural Initiative). We also thank Omar Davulcu at the Pacific Northwest Cryo-EM Center (PNCC, supported by NIH grant U24GM129547) for his help with data collection performed at the PNCC at OHSU and accessed through EMSL (grid.436923.9), a DOE Office of Science User Facility sponsored by the Office of Biological and Environmental Research. Research was supported by funding from the National Institutes of Health (R35 GM128786 to B.C., R01 DK107733 to E.B. and D.D.B, and R01 DK133229, R01 DK119360 to E.E.T.), the Welch Foundation (I-1944 to X.B.), and the National Science Foundation (CDF 2047640 to B.C. and D.D.B.).

## Author Contribution Statement

B.C. and E.B. conceived the project. E.B. oversaw cell biological and proteomic experiments performed by A.S. B.C. oversaw protein purification, biochemical experiments, and AlphaFold predictions performed by D.J.B. with the help from A.S.E., C.N., and D.A.K. Z.C. oversaw cryo-EM grid preparation, data collection, single particle reconstruction, and atomic-model building by B.C., Y.H., and H.Y.J.F. X.B. helped with cryo-EM data processing and supported cryo-EM grid preparation and screening performed by B.C. at UTSW. P.J. supervised cryo-EM grid preparation and data collection performed by D.J.B. at Iowa State. D.D.B. helped with cellular experiments and data interpretation. R.S. and E.E.T. performed cellular experiments related to LRMDA. B.C., H.Y.J.F., Z.C., and D.J.B. analyzed structures. E.B. and B.C. drafted the manuscript and prepared the figures with assistance from all other authors.

---

**Extended Data Fig. 1.**
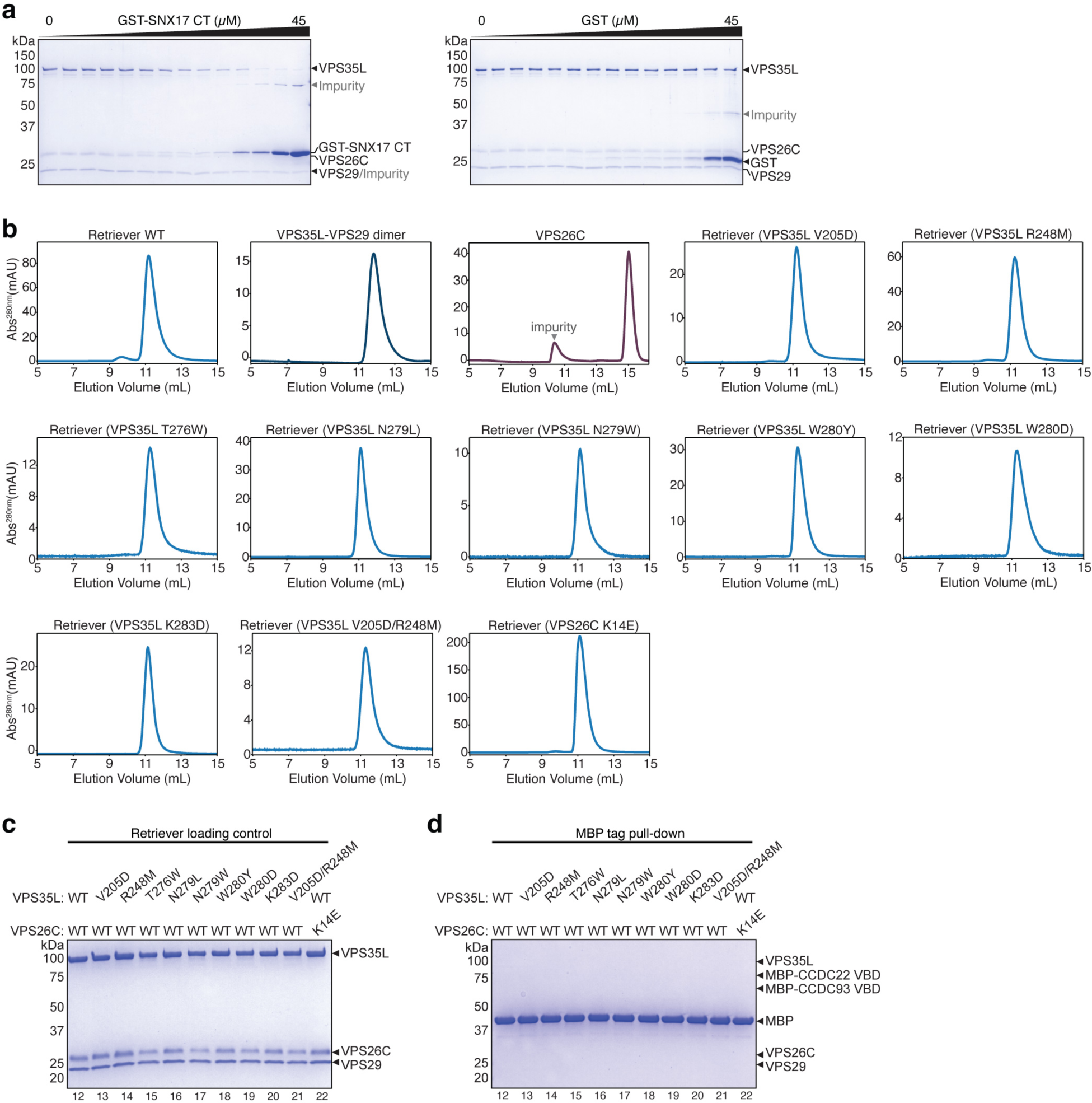
Quality control of proteins used in this study. **a.** Representative Coomassie blue-stained SDS PAGE gels showing the supernatant from the EPD assays used for quantification. **b-c.** Size exclusion chromatograms through a Superdex 200 Increase column **(b)** and Coomassie blue-stained SDS PAGE gels **(c)** showing loading control of indicated Retriever constructs used in pull-down assays shown in Fig. 2f. **d.** Coomassie blue-stained SDS PAGE gel showing MBP pull-down of indicated Retriever constructs using MBP-tag as a negative control, in comparison to MBP-CCDC22-CCDC93 VBD pull-down in Fig. 2g.

**Extended Data Fig. 2.**
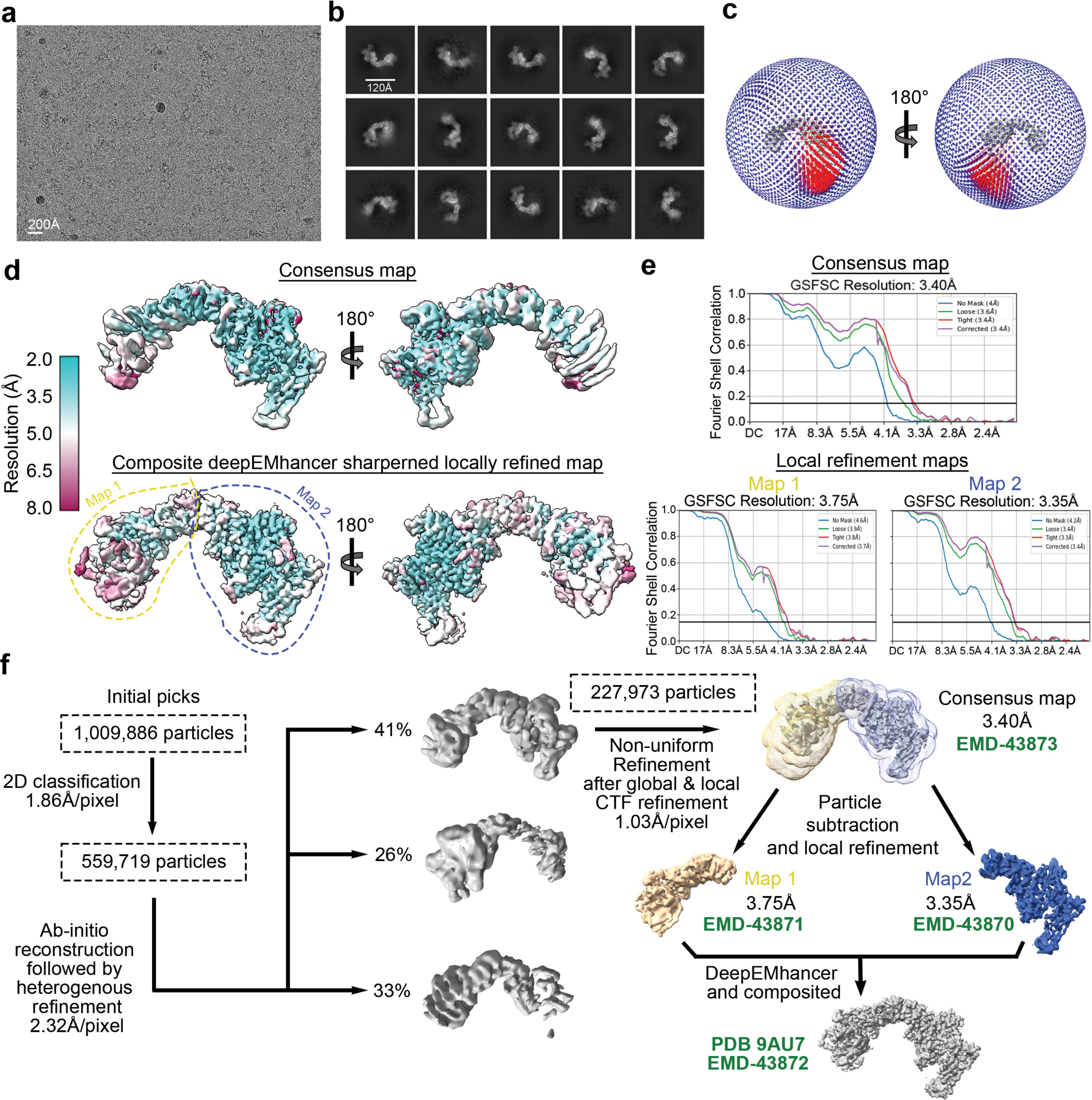
Cryo-EM data processing summary. **a.** Representative cryo-EM micrograph from a total of 10,009 micrographs used for data processing. **b.** Representative 2D class averages. **c.** 3D angular distribution of all Retriever-SNX17 particles that contributed to the final main map. **d.** Local resolution of the consensus map (upper) and the composite map stitched from two locally refined maps as indicated (lower). The orientation of the map is the same as in **(c)**. **e.** Fourier Shell Correlation (FSC) plots for the consensus map (upper) and the two locally refined maps (lower). **f.** Schematic showing cryo-EM data processing steps for obtaining 3D reconstruction of the Retriever-SNX17 complex. Maps and structural model deposited to PDB/EMDB are labeled.

**Extended Data Fig. 3.**
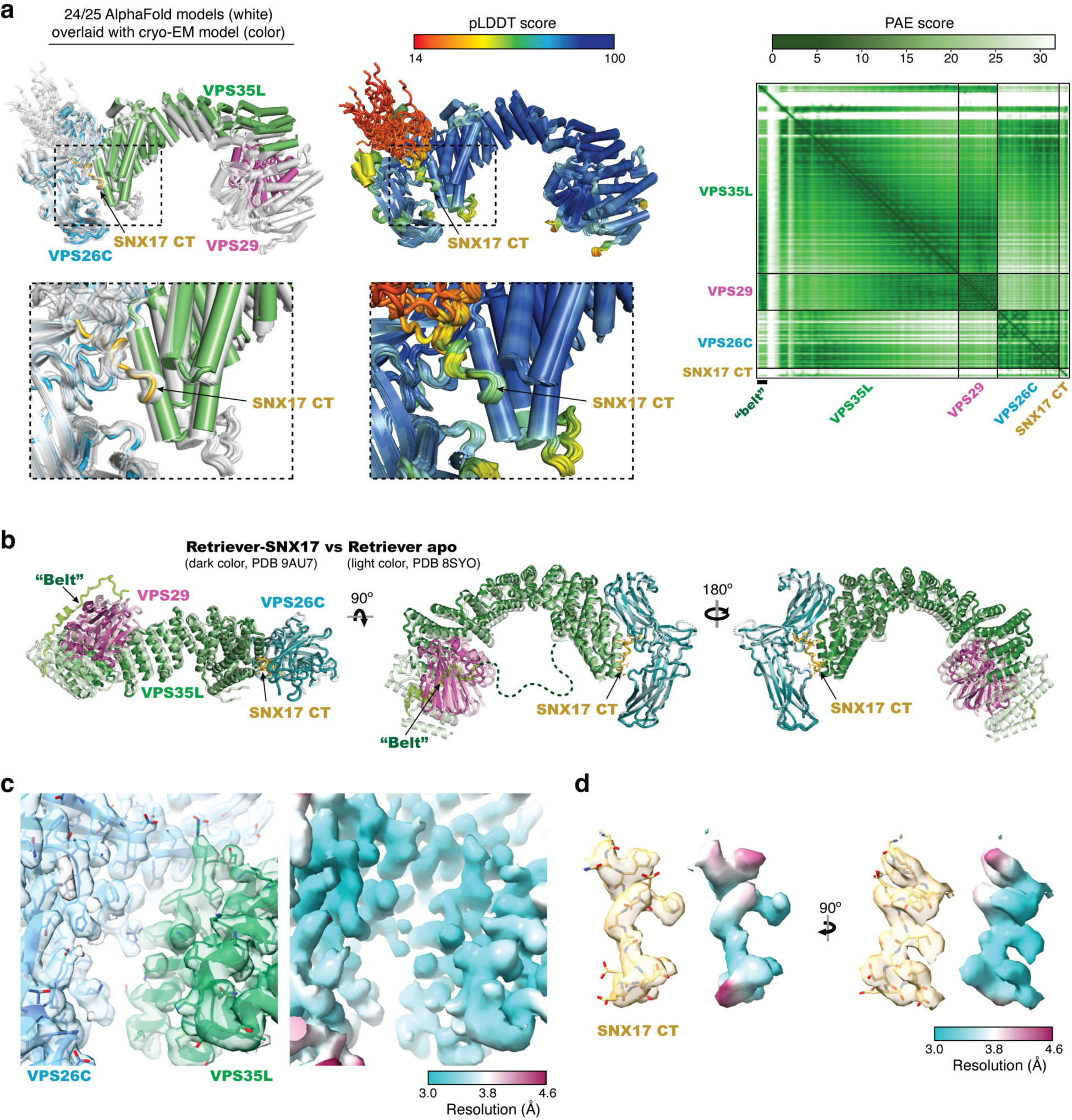
Overall structure comparison and local map quality. **a.** Left: Overlay of models derived from AlphaFold-Multimer prediction (white) and the cryo-EM model (color). Middle: Overlay of AlphaFold-Multimer models colored using predicted local difference distance test (pLDDT) scores for local structure accuracy. Right: Predicted aligned error (PAE) scores for distance error of the top-ranked AlphaFold-Multimer model. **b.** Overlay of the cryo-EM models between the Retriever-SNX17 complex (dark color, PDB 9AU7) and the Retriever apo form (light color, PDB 8SYO). **c-d.** Semi-transparent composite map overlayed with final model (left) (EMD-43872, PDB 9AU7) and the map colored by local resolution (right) showing the quality of the map and modeling of the SNX17 binding pocket in **(c)** and the SNX17 CT peptide in **(d)**.

**Extended Data Fig. 4.**
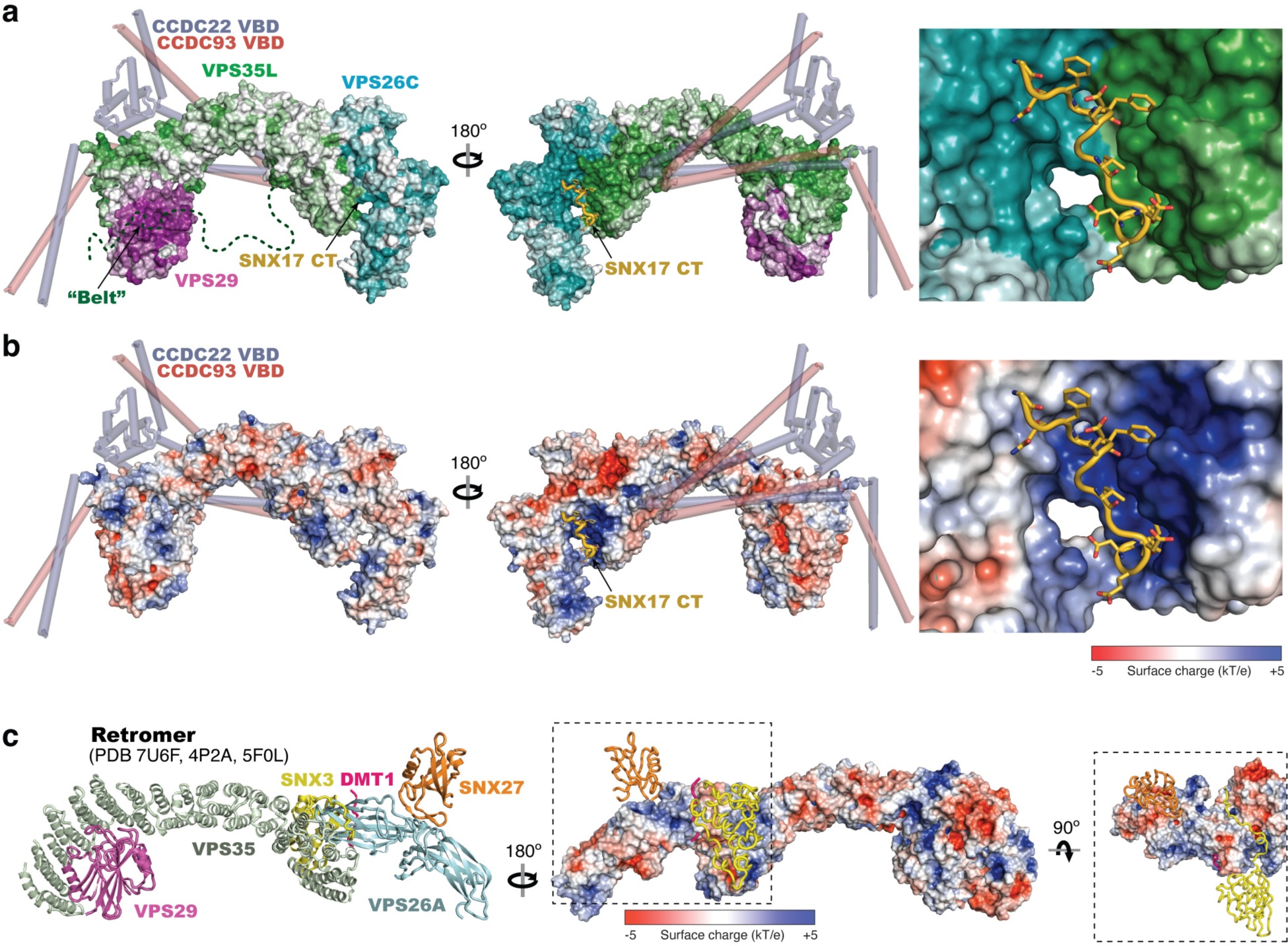
Surface conservation and electrostatic property. **a-b.** Overall surface conservation **(a)** and electrostatic potentials **(b)** of Retriever complexed with SNX17 CT peptide (shown in cartoon presentation). Surface conservation was calculated with ConSurf^33^, with color-to-white gradients representing the most (ConSurf score = 9) to the least conserved residues (ConSurf score = 1) following the same color scheme used in Fig. 2c, d. CCDC22-CCDC93 VBD model derived from AlphaFold-Multimer predictions are shown as semi-transparent cartoon presentation as a reference. **c.** Cartoon presentation (left) and surface presentation of electrostatic potentials (right) of Retromer, showing the binding sites for indicated ligands.

**Extended Data Fig. 5.**
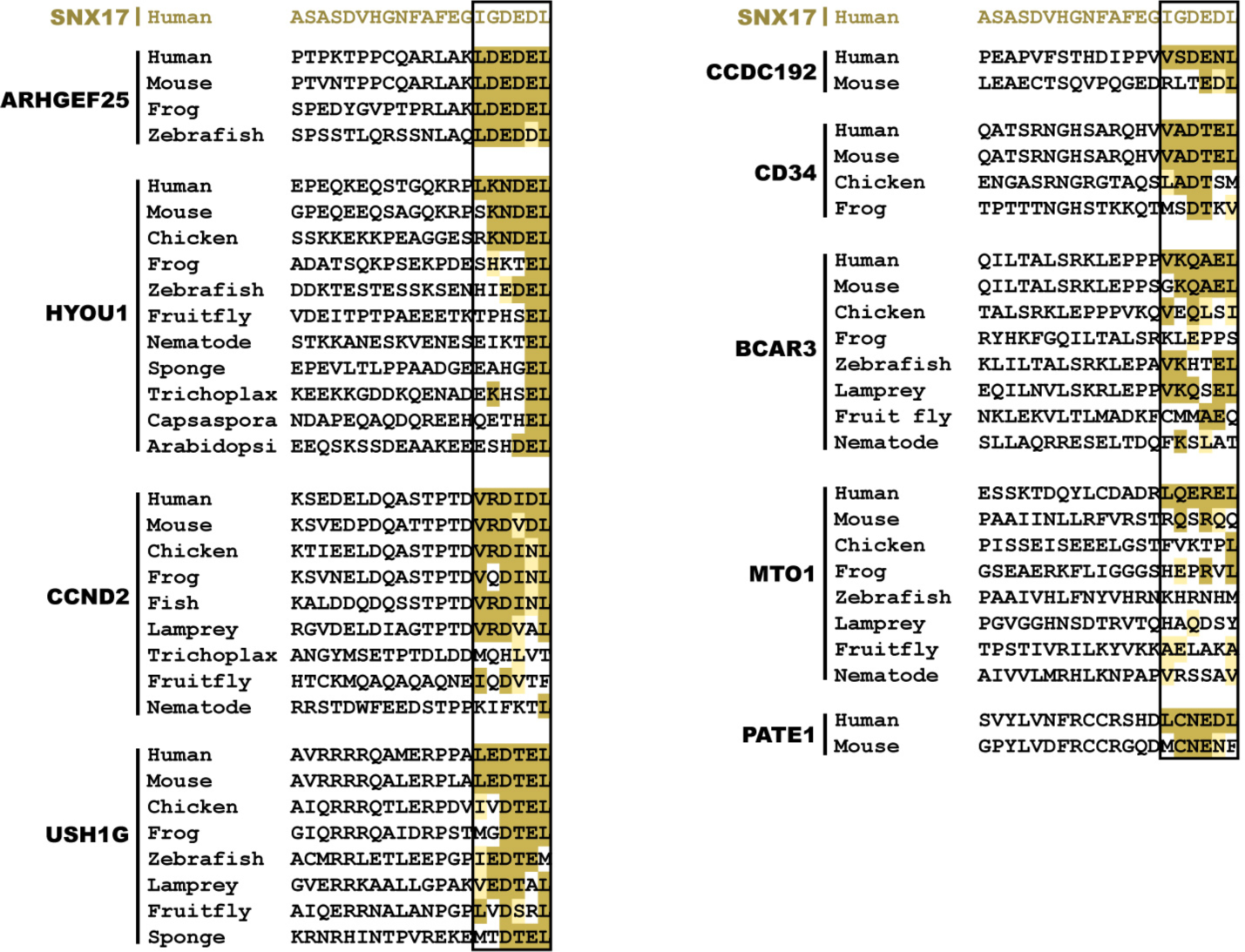
Sequence alignment of additional proteins potentially containing a SNX17 homology acidic tail. Gold, light-gold, and white shading denote sequences identical to, similar to, and not conserved with the human SNX17 CT sequence shown on the top, respectively. The 6-residue acidic sequences are highlighted by the black box.

## Methods

### Plasmids

All constructs were created using standard molecular biology procedures and verified by Sanger sequencing. Detailed information about constructs for recombinant protein production and mammalian expression, recombinant protein sequences, and DNA oligonucleotides for construct generation can be found in Supplementary Tables 1, 2, and 3, respectively. The ORFs of human VPS35L and VPS26C were previously described^34–36^. The mammalian expression vector for SNX17 was previously described^37^. SNX31 mammalian expression vector was designed by GeneArt at Thermo Fisher Scientific. For insect cell expression of Retriever, human full-length VPS35L (untagged, synthesized as a codon-optimized GeneString from Thermo Fisher to improve expression), VPS26C (untagged), and VPS29 (isoform 2) containing a C-terminal (GGS)_2_-His_6_ tag were cloned in a modified pFastBac^TM^ vector for insect cell expression as previously described^34^. For bacterial expression of isolated VPS26C, codon-optimized GeneString (Thermo Fisher) was cloned in a pMalC2Tev vector^34^. Bacterial expression vector of GST-SNX17 was previously described^38^. Bacterial expression vectors of CCDC22 VBD and CCDC93 VBD were previously described^34^. The CT 20 amino acids of SNX17, SNX31, LRMDA, TIMM23, PATE1, ARHGEF25, and HYOU1 were codon-optimized and cloned into a pGexTev vector using PCR.

### E. coli strains for protein expression

Standard, commercial *E. coli* strains used in this study include Mach1^T1R^ (Thermo Fisher) and BL21 (DE3)^T1R^ (Sigma), and are grown in Luria-Bertani medium using standard molecular biology conditions.

### Insect cell lines for protein expression

*Sf9* cells (Expression System) were maintained in Sf-900™ II serum-free medium (Thermo Fisher) and used for baculovirus preparation and large-scale expression.

### Cell culture

HEK293T (Cat # CRL-3216) and HeLa (Cat # CCL-2) cell lines were obtained from the American Type Culture Collection (Manassas, VA). Lenti-X 293T cells (Cat #632180) were obtained from Takara. All cell lines were cultured in high-glucose Dulbecco’s modified Eagle’s medium (DMEM) containing 10% fetal bovine serum (FBS) and 1% penicillin/streptomycin at 37°C with 5% CO_2_. Periodic PCR-based testing for Mycoplasma spp. was conducted to ensure culture purity. A HeLa line with VPS35L deficiency was previously described^39^ and these cells were complemented using a lentiviral vector to express HA tagged VPS35L protein versions as indicated.

### Transfection and lentiviral methods

HEK293T or Lenti-X 293T cells were transfected using Lipofectamine 2000 (Life Technologies) or PolyJet (SignaGen), respectively, and cultured for either 24 or 48 hours before analysis. VPS35L HeLa knockout cells were reconstituted with HA empty vector or various HA-tagged VPS35L using a lentivirus system. Lentivirus experiments followed a standard protocol as previously described for viral vector production and selection^40,41^.

### Immunofluorescence staining

We followed protocols previously described^36,39^. Briefly, cells were fixed with cold fixative (4% paraformaldehyde in PBS) for 18 min at room temperature in the dark, followed by 3-min permeabilization using 0.15% Surfact-Amps X-100 (28314, Thermo Fisher) in PBS. Samples were then incubated overnight at 4 °C in a humidified chamber with primary antibodies in immunofluorescence (IF) buffer (Tris-buffered saline plus human serum cocktail). After three washes in PBS, samples were incubated with secondary antibodies (1:500 dilution in IF buffer) for 1 h at room temperature or overnight at 4 °C in a humidified chamber. After four washes in PBS, coverslips were mounted on slides with SlowFade Anti-fade reagent (Life Technologies). Primary and secondary antibodies used are provided in Supplementary Table 4. Images were obtained using an A1R confocal microscope (Nikon, ×60 /1.4 oil immersion objective) operated by the NIS-Elements A1R (Nikon) software v5.42.03. Fluorescence signal values were quantified using Fiji v1.54f (ImageJ, NIH). Data were processed with Excel (Microsoft) and plotted with Prism v9.5.1 (GraphPad) or a Python web application https://biochempy.bb.iastate.edu. Each dot in the graphs represents the value from a single cell, with the horizontal bar indicating the mean and the error bars representing the standard error of the mean (SEM). Spearman’s Rank correlation coefficient was measured using EzColocalization Fiji Plugin within manually outlined regions of interest (ROIs).

### Mammalian protein extraction, immunoblotting, and immunoprecipitation

For most experiments, whole cell lysates were prepared using Triton X-100 lysis buffer (25 mM HEPES, 100 mM NaCl, 10 mM DTT, 1 mM EDTA, 10% Glycerol, 1% Triton X-100) supplemented with protease inhibitors (Roche). Immunoprecipitation, SDS-PAGE, and immunoblotting experiments were performed largely as previously described^39^. Specifically, for LRMDA immunoprecipitation, 48h after transfection, cells were harvested in NP-40 lysis buffer and mixed with anti-FLAG M2 magnetic beads (Sigma-Aldrich, Cat# F4799) for 2 hours at 4°C. The beads were washed 4 times in NP-40 lysis buffer, and the bound proteins were eluted with 150 μg/mL 3×Flag (Sigma-Aldrich, Cat# F4799) at 4 °C for 1.5 h. Western blot images were collected using ChemiDoc and Image Lab v6.1.0 (Biorad). Antibodies used are detailed in Supplementary Table 4.

### Cell surface biotinylation

Cell surface biotinylation was performed as previously reported^36^. Briefly, cells were incubated at 4⁰C with Sulfo-NHS-SS-biotin (Pierce) in biotinylation buffer (10 mM triethanolamine, 150 mM NaCl, 2 mM CaCl_2_, pH 8.0). After 30 min, cells were lysed in Tris-lysis buffer (50 mM Tris-HCl, pH 7.4, 150 mM NaCl, 1% NP-40, 0.5% Na deoxycholate, 5 mM EDTA, 5 mM EGTA) supplemented with Halt Protease/Phosphatase inhibitor (Thermo Fisher). Biotinylated proteins were captured using nanolink Streptavidin magnetic beads (Solulink) and washed three times with the same lysis buffer, once with high salt buffer (50 mM Tris-HCl, pH 7.4, 500 mM NaCl), and once with low salt buffer (10 mM Tris-HCl, pH 7.4, 5 μM Biotin). Proteins on the beads were eluted using at the elution buffer (PBS, 6M urea, 0.2% SDS (v/w) containing 100mM DTT at 65 °C for 30 min. For TMT proteomics, the eluted proteins were directly submitted in solution to the UT Southwestern Proteomics core facility.

### Protein affinity purification

Knockout cells expressing HA-tagged VPS35L were grown on culture dishes and lysed in Triton-X lysis buffer. Clarified cell lysates containing equal amounts of protein were added to HA-resin to capture HA-tagged proteins. HA beads were washed using lysis buffer and eluted using 3 x LDS/DTT gel loading buffer at 95 °C. Eluted proteins were analyzed by SDS-PAGE and LC-MS/MS mass spectrometry at the UT Southwestern Proteomics core.

### Proteomic interactome and cell surface analysis

We combined protein identification, abundance (based on spectral index), and enrichment ratios (compared to empty vector) to identify potential interacting proteins. After reduction with DTT and alkylation with iodoacetamide (Sigma–Aldrich), samples were digested overnight with trypsin (Pierce). After solid-phase extraction cleanup with an Oasis HLB plate (Waters), digested samples were injected into an Orbitrap Fusion Lumos mass spectrometer coupled to an Ultimate 3000 RSLC-Nano liquid chromatography system. Through a 75 μm i.d., 75-cm long EasySpray column (Thermo), samples were eluted with a gradient from 1-28% buffer B over 90 min. Buffer A contained 2% (v/v) ACN and 0.1% formic acid in water, and buffer B contained 80% (v/v) ACN, 10% (v/v) trifluoroethanol, and 0.1% formic acid in water. The mass spectrometer operated in positive ion mode with a source voltage of 1.8-2.4 kV and an ion transfer tube temperature of 275 °C. MS scans were acquired at 120,000 resolution in the Orbitrap. Uto 10 MS/MS spectra were obtained in the ion trap for each full spectrum acquired using higher-energy collisional dissociation (HCD) for ions with charges 2-7. Dynamic exclusion was set for 25 s after an ion was selected for fragmentation. For the plasma membrane and interaction proteomics samples, raw MS data were analyzed using Proteome Discoverer v3.0 (Thermo), with peptide identification performed using Sequest HT searching against the human protein database from UniProt. We set fragment and precursor tolerances at 10 ppm and 0.6 Da, respectively, and allowed three missed cleavages. We set cysteine carbamidomethylation as a fixed peptide modification and methionine oxidation as a variable modification. We applied a false-discovery rate (FDR) cutoff of 1% for all peptides.

To analyze protein complex composition in native gel samples, raw MS data were analyzed using MaxQuant v.2.0.3.0, with peptide identification performed against the human protein database from UniProt. We set fragment and precursor tolerances at 20 ppm and 0.5 Da, respectively, and allowed three missed cleavages. We set cysteine carbamidomethylation as a fixed peptide modification, and methionine oxidation and N-terminal acetylation as a variable modification. We used iBAQ quantitation for protein quantitation within each sample.

### TMT proteomics

For TMT-based proteomic quantification, samples were thoroughly mixed with 25 μL of 10% SDS and 100 mM triethylammonium bicarbonate (TEAB) by vortexing and then reduced by 2 μL of 0.5 M tris(2-carboxyethyl)phosphine (TCEP) at 56 °C for 30 min. Free cysteines were then alkylated by 2 μL of 500 mM iodoacetamide in the dark at room temperature for 30 min. Afterwards, samples were added with 5.4 μL of 12% phosphoric acid and 300 μL of S-Trap (Protifi) binding buffer before being loaded onto an S-Trap column. Samples were digested by 1 μg of trypsin overnight at 37 °C. Digested peptides were dried and reconstituted in 21 μL of 50 mM TEAB buffer. Based on absorbance at 205 nm using NanoDrop, equal amounts of peptides were labelled with TMT 6plex reagent (Thermo), quenched with 5% hydroxylamine, combined, dried in a SpeedVac, desalted using an Oasis HLB microelution plate (Waters), and dried again in a SpeedVac. Finally, samples were dissolved in 50 μL of 2% acetonitrile and 0.1% TFA and then injected onto an Orbitrap Eclipse mass spectrometer coupled to an Ultimate 3000 RSLC-Nano liquid chromatography system. Samples were developed through a 75 μm i.d., 75-cm long EasySpray column (Thermo) and eluted with a gradient from 1-28% buffer B over 180 min, followed by 28-45% buffer B over 25 minutes. Buffer A contained 2% (v/v) ACN and 0.1% formic acid in water, and buffer B contained 80% (v/v) ACN, 10% (v/v) trifluoroethanol, and 0.1% formic acid in water. The mass spectrometer operated in positive ion mode with a source voltage of 2.0 kV and an ion transfer tube temperature of 300 °C. MS scans were acquired at 120,000 resolution in the Orbitrap over a mass range of m/z = 400-1600, and top speed mode was used for SPS-MS3 analysis with a cycle time of 2.5 s. MS2 was performed using collisionally-induced dissociation (CID) with a collision energy of 35% for ions with charges 2-6. Dynamic exclusion was set for 25 s after an ion was selected for fragmentation. Real-time search was performed using the reviewed human protein database from UniProt. We set cysteine carbamidomethylation and TMT 6plex modification of lysine and peptide N-termini as fixed modifications, and methionine oxidation as a variable modification. We allowed two missed cleavages and up to 3 modifications per peptide. The top 10 fragments for MS/MS spectra corresponding to peptides identified by real-time search were selected for MS3 fragmentation using high-energy collisional dissociation (HCD), with a collision energy of 65%. Raw MS data files were analyzed using both the Sequest HT and Comet nodes within Proteome Discoverer v3.0 (Thermo), searching against the reviewed human protein database from UniProt. Fragment and precursor tolerances of 10 ppm and 0.6 Da were specified, and two missed cleavages were allowed. The same modifications were used in the search as for the real-time search. The false-discovery rate (FDR) cutoff was 1% for all peptides.

### Recombinant protein purification

The Retriever complex or the VPS35L-VPS29 subcomplex was expressed from *Sf9* cells using the Bac-to-Bac system and purified through Ni-NTA affinity, cation exchange, and anion exchange, and size exclusion chromatography essentially as previously described^34^. To improve expression, VPS35L ORF was changed to a codon-optimized sequence from Thermo Fisher. Typical yield was ∼1 mg of purified Retriever from 3 liters of *Sf9* culture. SNX17, isolated VPS26C, and SNX17-homologous CT tails were expressed and purified following procedures essentially as previously described^34^. Briefly, proteins were expressed in BL21 (DE3)^T1R^ cells (Sigma) at 18 °C overnight after induction with 1 mM IPTG. GST-tagged SNX17 proteins were purified using Glutathione Sepharose beads (Cytiva) and eluted using 100 mM Tris pH 8.5, 50 mM NaCl, and 30 mM reduced glutathione. The resulting GST-SNX17 proteins were further purified by anion exchange chromatography using a 2-mL Source 15Q column (10 mM Tris pH 8.0 and 5 mM BME in a gradient of 0 - 400 mM NaCl developed over 40 mL) and size exclusion chromatography using a 24-mL Superdex Increase 200 column [10 mM HEPES pH 7.0, 100 mM NaCl, 5% (w/v) glycerol, and 1 mM DTT]. SNX17 point mutants were purified using Glutathione Sepharose beads as described above and then dialyzed into 10 mM HEPES pH 7.0, 100 mM NaCl, 5% (w/v) glycerol, and 1 mM DTT for pull-down assays. SNX17-homologous C-terminal tails were purified using Glutathione Sepharose beads as described above, further purified by a 2-mL Source 15Q column (10 mM Tris pH 8.0 and 5 mM BME in a gradient of 0 - 400 mM NaCl developed over 40 mL), and finally dialyzed into 10 mM HEPES pH 7.0, 50 mM NaCl, 5% (w/v) glycerol, and 1 mM DTT for pull-down assays. Isolated MBP-tagged VPS26C was purified using Amylose beads (New England Biolabs) and eluted using 20 mM Tris pH 8.0, 200 mM NaCl, 2% (w/v) maltose, and 5 mM BME. The protein was further purified by anion exchange chromatography using a 2-mL Source 15S column (10 mM HEPES pH 7.0 and 5 mM BME in a gradient of 0 - 400 mM NaCl developed over 40 mL) and cleaved using TEV protease to remove the MBP tag. Cleaved VPS26C was polished by size exclusion chromatography using a 24-mL Superdex Increase 75 column [10 mM HEPES pH 7.0, 100 mM NaCl, 5% (w/v) glycerol, and 1 mM DTT]. MBP-CCDC22 VBD and MBP-CCDC93 VBD were purified as described^34^. All chromatography steps were performed using Cytiva columns on an ÄKTA^TM^ Pure protein purification system. SNX17 C-terminal peptide, corresponding to amino acid sequence 451-470, ASASDVHGNFAFEGIGDEDL, was synthesized from GenScript at ≥98% purity. The lyophilized peptide was dissolved in 100 mM HEPES pH 7.0 buffer at a stock concentration of 40 mg/mL (19.5 mM), aliquoted in small volumes, and stored at −80 °C.

### In vitro pull-down assays

GST pull-down experiments followed previous procedures^42^. Briefly, bait (100-200 pmol of GST-tagged proteins) and prey (50-200 pmol for Retriever) were mixed with 20 µL of Glutathione Sepharose beads (Cytiva) in 1 mL of binding buffer [10 mM HEPES pH 7, 50 mM NaCl, 5% (w/v) glycerol, 0.05% (w/v) Triton-X100, and 5 mM BME] at 4 °C for 30 min. After three 1-mL washes with the binding buffer, bound proteins were eluted with 100 mM Tris pH 8.5, 50 mM NaCl, and 30 mM reduced glutathione and examined by SDS-PAGE. Where it is indicated, 200 pmol of MBP-CCDC22-CCDC93 VBD dimer or various amounts of SNX17 CT peptide were added in pull-down assays.

### In vitro equilibrium pull-down (EPD) assays

Equilibrium pull-down assays were performed as previously described^42^. Briefly, 60 µL of Glutathione Sepharose beads (50% slurry equilibrated in a pull-down buffer [10 mM HEPES pH 7, 50 mM NaCl, 5% (w/v) glycerol, 0.05% (w/v) Triton-X100, and 5 mM BME] were mixed with 0.1 µM Retriever and various amounts of GST-tagged protein (up to 45 µM, stored in the same pull-down buffer) and brought to 100 µL final reaction volume using the pull-down buffer. The reactions were allowed to mix for 30 min at 4 °C, and four reactions at a time were spun at 15 krpm for 15 seconds. The supernatant was immediately removed and examined by SDS-PAGE and Coomassie blue staining. The VPS35L intensity was quantified using ImageJ v2.3.0/1.53q to calculate the fractional occupancy. The data from all repeats were pooled and globally fitted in DynaFit v4.08.187 using a single binding site model^43,44^.

### Sample preparation for electron microscopy

Purified Retriever and synthesized SNX17 CT peptide were mixed freshly at a final concentration of 1.4 µM Retriever and 6.5 mM peptide in a final buffer containing 10 mM HEPES pH 7.0, 150 mM NaCl, 5% (w/v) glycerol, and 1 mM DTT. The mixture was centrifuged for at least 10 min at 4 °C before 3 µL was applied to a glow-discharged Quantifoil 300-mesh R1.2/1.3 Copper grid (Micro Tools GmbH). After a 10-second preincubation under 100% humidity at 4°C, the grid was blotted for 3.5 sec and plunge-frozen in liquid ethane using Vitrobot Mark IV (Thermo Fisher).

### Electron microscopy data acquisition

Sample grids were screened on a 200 kV Talos Artica or Glacios microscope (Thermo Fisher) at the Cryo Electron Microscopy Facility by Structural Biology Laboratory at University of Texas Southwestern Medical Center (UTSW) or at the cryo-EM Facility at Iowa State University. The final cryo-EM data were acquired on a Titan Krios microscope (Thermo Fisher) at PNCC operated at 300 kV, with a post-column energy filter (Gatan) and a K3 direct detection camera (Gatan) in non-CDS mode. Movies were acquired using SerialEM v4.0^45^ at a pixel size of 0.4133 Å in super-resolution counting mode, with an accumulated total dose of 60 e^−^/Å^2^ over 60 frames. The defocus range of the images was set between −0.9 to −2.5 μm. In total, 10,009 movies were collected for data processing.

### Electron Microscopy data processing

Cryo-EM data were processed using cryoSPARC^46^ v4.4.1. Pre-processing was performed in cryoSPARC Live, including motion correction with a binning factor of 2, resulting in a pixel size of 0.8266 Å/pixel and Contrast Transfer Function (CTF) estimation. Blob picking was used in cryoSPARC Live and 1,009,886 particles were selected after initial 2D classification. After extensive 2D classification, 559,719 particles were selected for *ab initio* 3D reconstruction and heterogeneous refinement (Extended Data Fig. 2). The best resolved 3D class, containing 227,973 particles, underwent global and local CTF-refinement and a final non-uniform refinement, producing a full map with an overall resolution of 3.4 Å with a binned pixel size of 1.03 Å/pixel (deposited in EMD-43873 with its associated half maps). To improve the map quality on both ends of the map, two masks were generated—one for VPS29-bound half of VPS35L, and another for VPS26C-bound half. Signals outside the masks were subtracted and local refinement of the two regions were performed independently. The resulted maps were deposited with their half maps at EMD-43871 and EMD-43870, respectively (Extended Data Fig. 2d-f). DeepEMhancer v20220530_cu11^47^ was then used with the unfiltered half maps to generate sharpened maps of the two locally refined maps, respectively, and a composite of the two was generated in UCSF ChimeraX v1.6.1 by the vop maximum command^48^. This composite map (EMD-43872/PDB-9AU7) was used for modeling and shown in Fig. 2 and Extended Data Fig 3c, d. All reported resolutions followed the gold-standard Fourier shell correlation (FSC) using the 0.143 criterion^49^.

### Atomic model building

A model of the Retreiver-SNX17 complex predicted by AlphaFold Multimer v2.3.1 was used as the initial model^50^ for model building. Model was first docked and fitted into the sharpened composite map using ISOLDE^51^ in ChimeraX, followed by iterations of real-space refinement in Phenix v1.21^52^ with reference model and secondary structure restraints and manual building in COOT v0.9.8.8^53,54^. Model geometries were assessed by MolProbity in Phenix^55^ (http://molprobity.biochem.duke.edu/), and the PDB Validation server^56^ (www.wwpdb.org). Figures were generated using PyMOL v2.5.4 or ChimeraX v1.7.1^57^.

### AlphaFold prediction and analysis

AlphaFold version 2.3.1 (https://github.com/deepmind/alphafold) was installed on local NVidia A100 80GB GPU computers at Iowa State University ResearchIT or High-Performance Computing for AlphaFold Multimer prediction using standard AlphaFold procedures^50,53^ as previously described^34^.

### Reproducibility and statistical analysis

To assess statistical significance, one-way ANOVA with Dunnett’s post-hoc test was applied to compare multiple groups with one control group, using Prism v9.5.1 (GraphPad). An error probability below 5% (p < 0.05; * in Figure panels) was considered to imply statistical significance. All imaging and co-precipitation experiments were performed in two to four independent iterations. All in vitro pull-down assays were performed at least twice, unless otherwise indicated. Large scale proteomics were performed once, with key results confirmed using other methods.

## Data availability

Cryo-EM maps and models have been deposited in the EMDB (accession number EMD-43870, EMD-43871, EMD-43872, and EMD-43873) and PDB (accession number 9AU7). AlphaFold Multimer-derived models are available in ModelArchive (modelarchive.org) with the accession code ma-swt4h. Mass spectrometry data have been deposited at the MassIVE repository (accession numbers MSV000094100 and MSV000094101). Source data are available for all uncropped western blots, Coomassie-blue gels, and all quantitative datasets presented here. To our knowledge, all information required to reanalyze the data reported here is publicly available. Any additional data we inadvertently missed will be shared upon reasonable request. This paper does not report original code.

---

**Supplementary Table 1:**
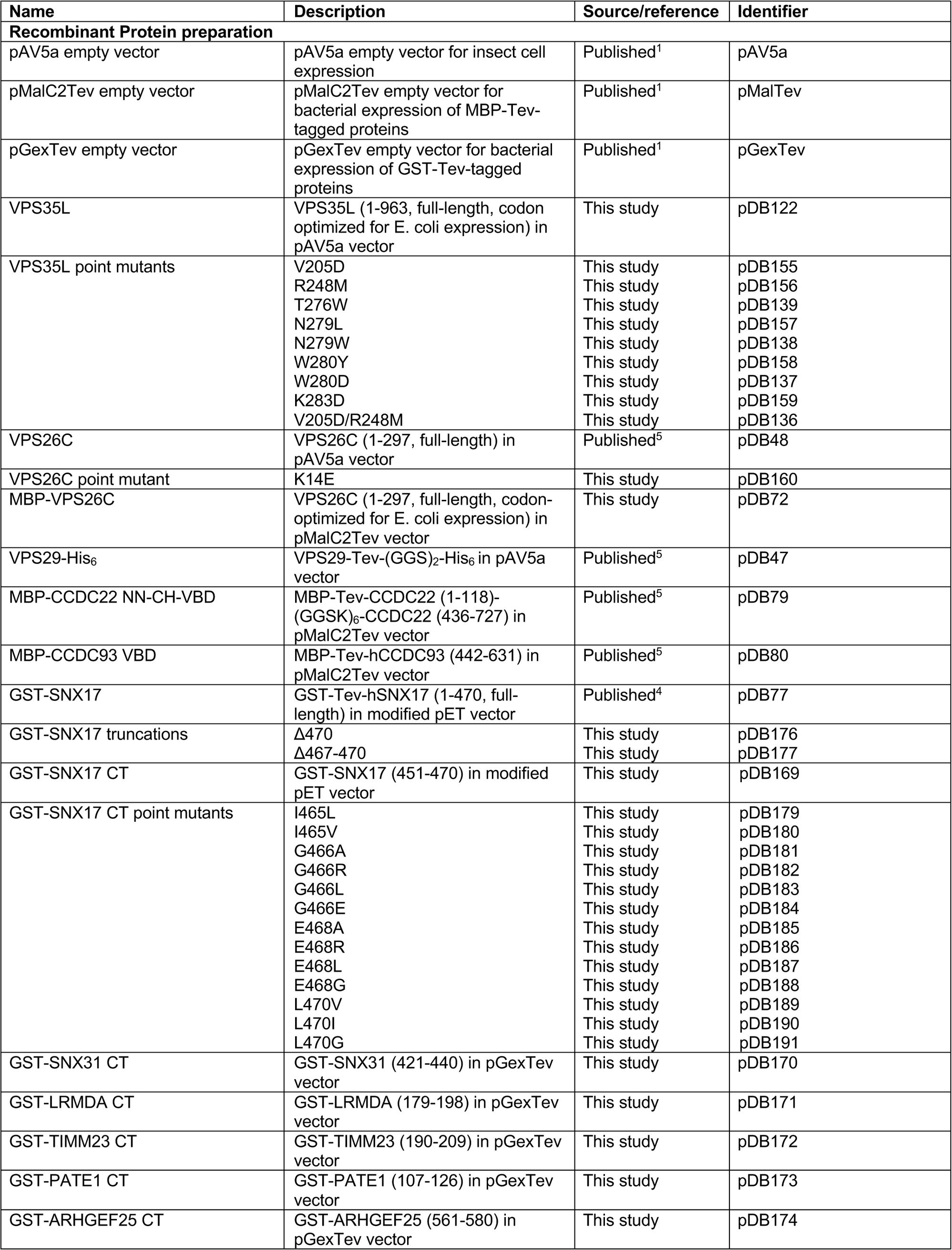
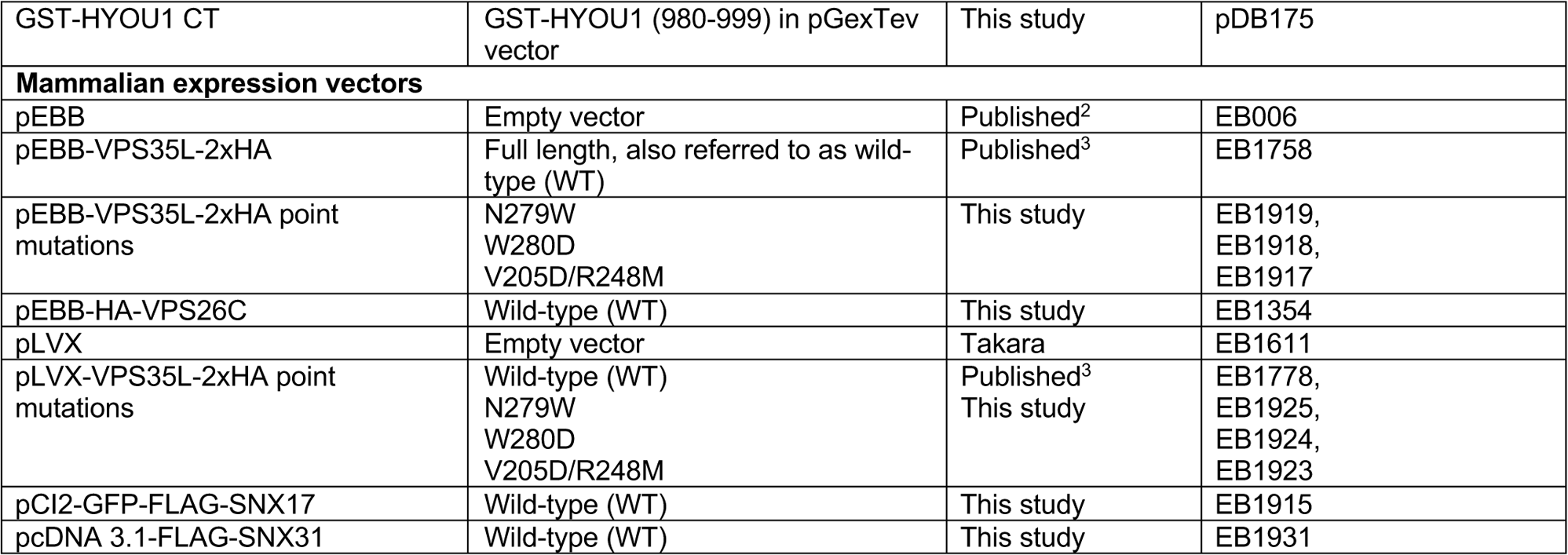
DNA constructs used in this study.

**Supplementary Table 2:**
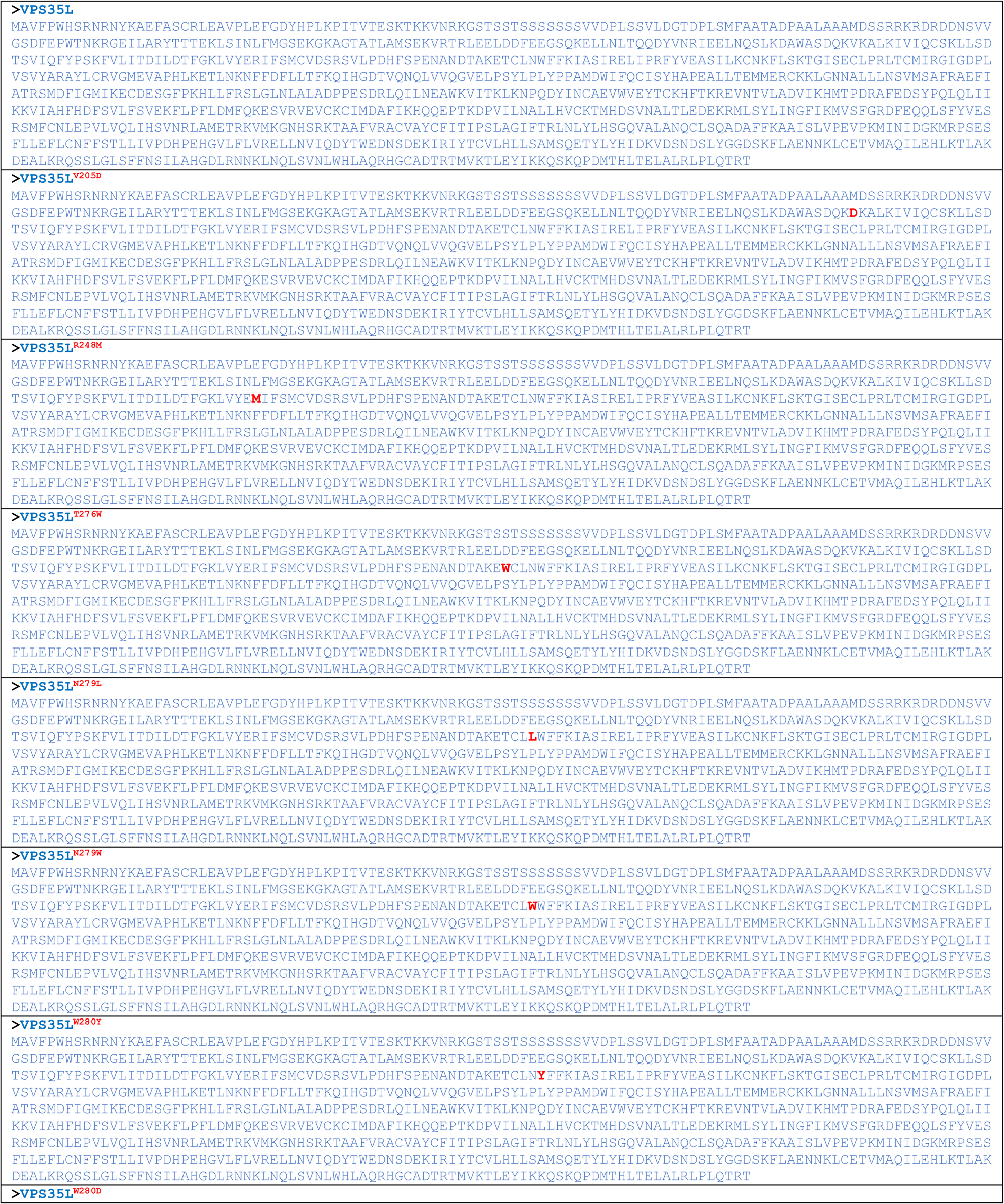
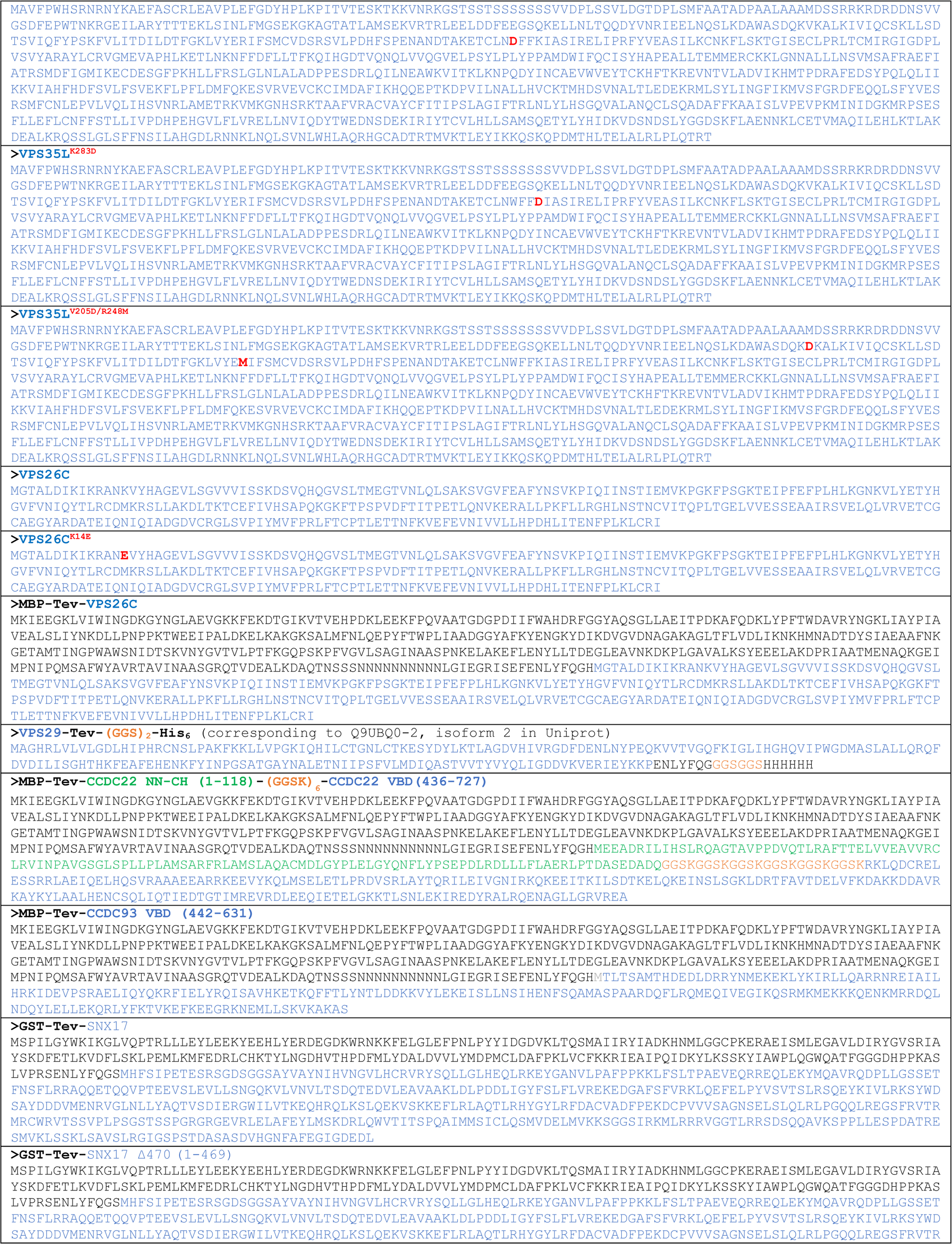
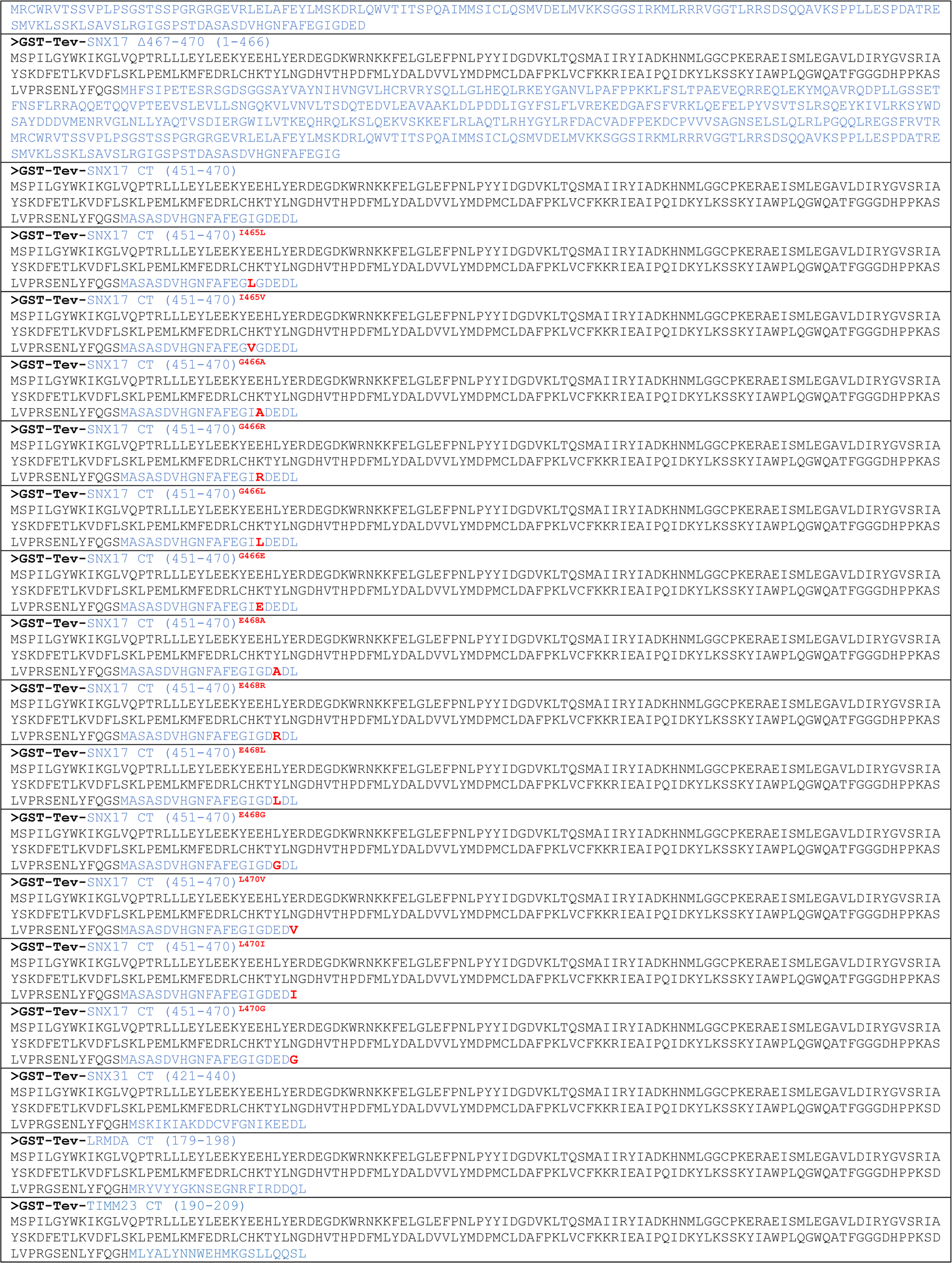
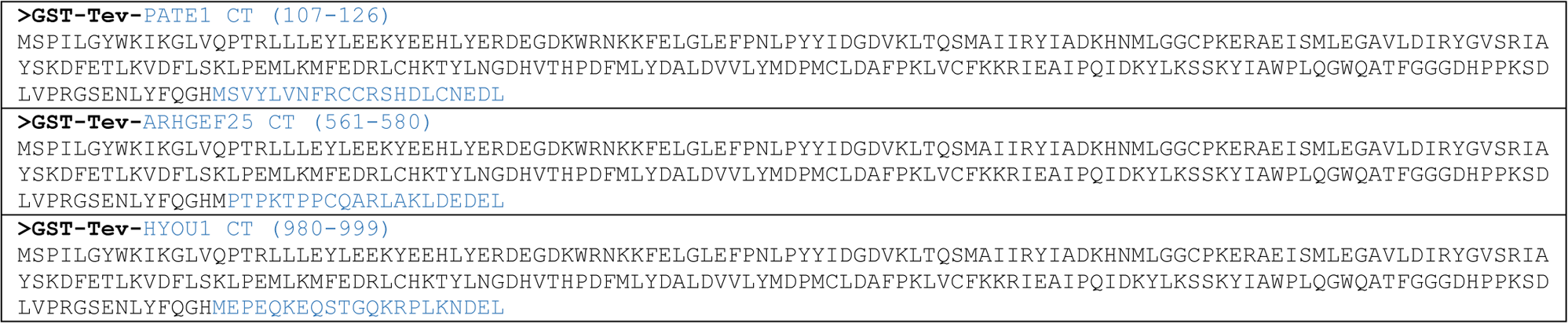
Sequences of recombinant proteins used in this study. Only sequences in the final product (i.e., after protease cleavage to remove the affinity tag) are shown and are annotated by corresponding colors.

**Supplementary Table 3:**
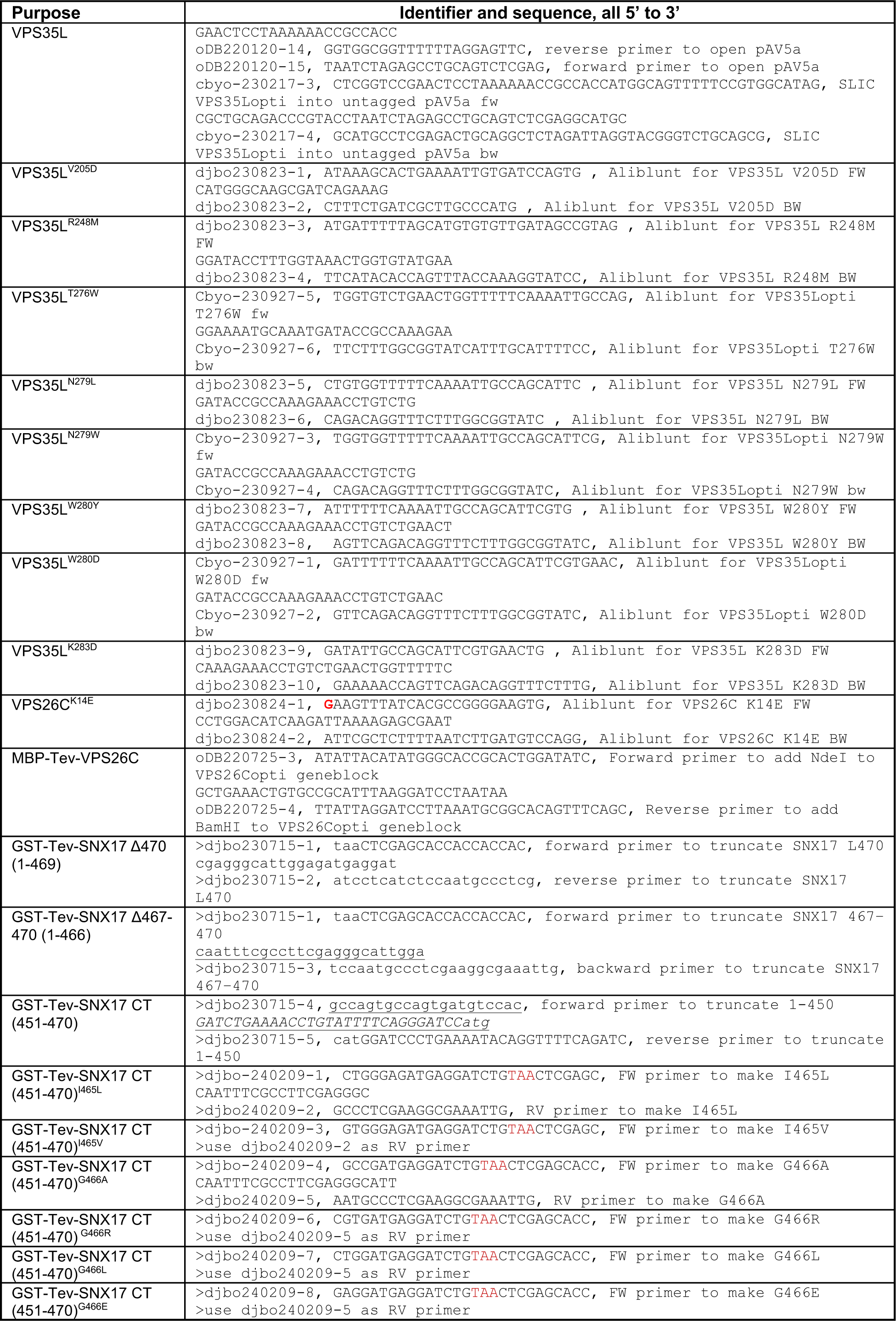
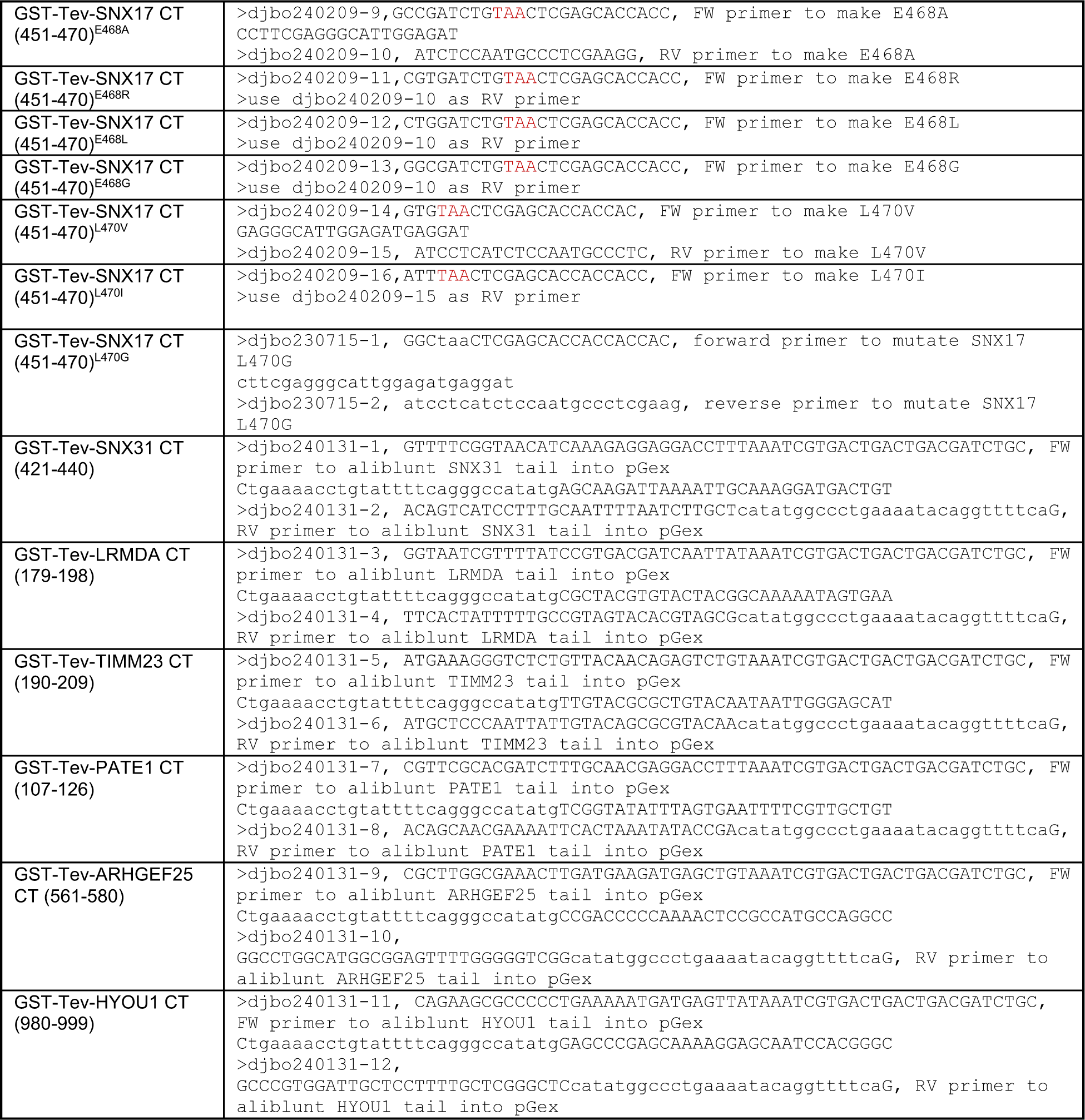
DNA oligos used in this study.

**Supplementary Table 4:** Antibodies used in this study.

| Primary antibodies |  |  |  |  |
| --- | --- | --- | --- | --- |
| WB, western blot; IF, immunofluorescence staining; FC, Flow Cytometry. |  |  |  |  |
| Target | Source (host species) | Catalog, Clone (References) | Application (dilution) | Validation information |
| CCDC22 | ProteinTech Group (rabbit) | 16636-1-AP | WB (1:1000) | Phillips-Krawczak et al., 2015<sup>4</sup> |
| CCDC93 | ProteinTech Group (rabbit) | 20861-1-AP | WB (1:1000) | Phillips-Krawczak et al., 2015<sup>4</sup> |
| COMMD1 | ProteinTech Group (rabbit) | 11938-1-AP | WB (1:1000) | Manufacturer validation using various tissues and IF |
| DENND10 | Custom made, Cocalico Biologicals (rabbit) | 95-110 (Singla et al., 2019<sup>6</sup>) | WB (1:500) | Singla et al., 2019<sup>6</sup> |
| FAM21 | Custom made, Cocalico Biologicals (rabbit) | MC2188 (Gomez and Billadeau, 2009<sup>7</sup>) | IF (1:1000) | Gomez and Billadeau, 2009<sup>7</sup> |
| FLAG | Sigma (mouse) | F1804, M2 | WB (1:500) | Manufacturer validation with overexpressed proteins |
| HA | Biolegend (mouse) | 901502, 16B12 | WB (1:500), IF (1:100) | Manufacturer validation with overexpressed proteins |
| HA | Cell Signaling (mouse) | 2999S, 6E2 | WB (1:1000) | Manufacturer validation with overexpressed proteins |
| Integrin-α5 | BD Biosceinces (mouse) | 555615, IIA1 | IF (1:100) | Manufacturer validation in FACS using isotype control |
| Integrin-β1 | Santa Cruz (mouse) | sc-53711, TS2/16 | IF (1:100) | Manufacturer validation in FACS using isotype control |
| LRMDA | Abcam (rabbit) | 150986 | WB (1:500) | Manufacturer validation with overexpressed proteins |
| TIMM23 | Proteintech Group (rabbit) | 11123-1-AP | WB (1:500) | Manufacturer validation with immunoprecipitation |
| VPS26C | Millipore (rabbit) | ABN87 | WB (1:5000) | Singla et al., 2019<sup>6</sup> |
| VPS29 | GeneTex (rabbit) | GTX104768 | WB (1:500) | Singla et al., 2019<sup>6</sup> |
| Secondary antibodies used for immunofluorescence |  |  |  |  |
| Fluorophore | Source (target species) | Catalog number (dilution) |  | Validation |
| Alexa 488 | Invitrogen (mouse) | A11029 (1:500) |  | Manufacturer validation in IF using no primary antibody controls |
| Alexa 555 | Invitrogen (rabbit) | A21428 (1:500) |  |  |

